# Multi-model functionalization of disease-associated *PTEN* missense mutations identifies multiple molecular mechanisms underlying protein dysfunction

**DOI:** 10.1101/800011

**Authors:** Kathryn L. Post, Manuel Belmadani, Payel Ganguly, Fabian Meili, Riki Dingwall, Troy A. McDiarmid, Warren M. Meyers, Caitlin Herrington, Barry P. Young, Daniel B. Callaghan, Sanja Rogic, Matthew Edwards, Ana Niciforovic, Alessandro Cau, Catharine H. Rankin, Timothy P. O’Connor, Shernaz X. Bamji, Christopher J. Loewen, Douglas W. Allan, Paul Pavlidis, Kurt Haas

## Abstract

Functional variomics provides the foundation for personalized medicine by linking genetic variation to disease expression, outcome and treatment, yet its utility is dependent on appropriate assays to evaluate mutation impact on protein function. To fully assess the effects of 106 missense and nonsense variants of PTEN associated with autism spectrum disorder, somatic cancer and PHTS, we take a deep phenotypic profiling approach using 18 assays in 5 model systems spanning diverse cellular environments ranging from molecular function to neuronal morphogenesis and behavior. Variants inducing instability occurred across the protein, resulting in partial to complete loss of function (LoF), which was well correlated across models. However, assays were selectively sensitive to variants located in substrate binding and catalytic domains, which exhibited complete LoF or dominant negativity independent of effects on stability. Our results indicate that full characterization of variant impact requires assays sensitive to instability and a range of protein functions.

## INTRODUCTION

Autism spectrum disorder (ASD) is the most common genetic neurodevelopmental disorder, characterized by impaired social communication and restricted, repetitive behaviors^1^. Headway in understanding ASD pathophysiology has been stymied by tremendous phenotypic heterogeneity and complex genetics across individuals. Conclusively linking genes to ASD is challenging since sequencing efforts have identified mutations in hundreds of genes from individuals with the disorder^2–9^. Confidence of gene association to ASD is enhanced by identification of *de novo,* likely gene disrupting (LGD) mutations; however, the most common *de novo* mutations identified in individuals with ASD are missense (MS) variants of unknown significance (VUS)^2,9^. A complete understanding of ASD genetics and their link to pathophysiology requires deciphering how these mutations impact protein function, and how protein dysfunction affects neural circuits and behavior^10^.

The lipid and protein phosphatase, PTEN (phosphatase and tensin homologue deleted on chromosome 10), has well-established links to ASD^11^, somatic cancers^12^, and the cancer-predisposing PTEN hamartoma syndrome (PHTS)^13,14^. However, while LGD mutations in *PTEN* have been found in individuals with ASD, the majority of variants identified are MS VUS, and their impact on protein function and causal links to the molecular mechanism(s) of disease expression remain unclear. Understanding the specific impact of these point mutations on PTEN function and their repercussions in cellular and tissue development is necessary for relating its disruption to disease.

As PTEN is a multifunctional protein expressed throughout the body, and both cancer and ASD are complex conditions involving diverse tissues, it is likely that VUS may disrupt distinct PTEN functions in different cell types and developmental conditions to contribute to disease. Saturation mutagenesis and massively parallel functional approaches to characterize single nucleotide variants of PTEN have identified a large range of impacts on PTEN’s lipid phosphatase activity using single-cell assays^15,16^. In order to more comprehensively assess a full range of PTEN functions, here we take a novel deep phenotypic profiling approach by assessing the impact of MS and nonsense (NS) variants in 18 bioassays in 5 model systems. Applying high-throughput unbiased analyses of genetic interactions of human PTEN in *Saccharomyces cerevisiae* we develop a novel assay of PTEN variant function sensitive to lipid phosphatase activity. By expressing human *PTEN* variants in *Drosophila melanogaster* we assess the rate of development in an invertebrate model, regulated by the insulin receptor pathways^17,18^. We test a smaller subset of *PTEN* variants for effects on neuronal dendritic and axonal growth, and excitatory and inhibitory synaptogenesis using rat primary neuronal cultures, processes found to be disrupted in ASD models^19,20^ and mediated by both lipid and protein phosphatase PTEN activities^21,22^. As ASD is primarily defined as disordered behavior and sensorimotor processing, we test impact of variants on chemotaxis in *Caenorhabditis elegans*. In order to probe the molecular mechanisms causing variant dysfunction in these assays, we performed flow cytometry analysis of human embryonic kidney (HEK293) cells overexpressing *PTEN* variants to parse effects on protein stability and catalytic activity. These multi-model results provide robust validation of measures of variant function in highly diverse cellular environments to uncover complexities of the contribution of single nucleotide mutations to protein dysfunction and pathophysiology.

## RESULTS

### Categorization of PTEN variants

We selected PTEN MS and NS mutations identified in individuals with ASD, intellectual disability (ID), developmental delay (DD), somatic cancer and PHTS, as well as variants found among the general population (**Fig 1a, Supp. Table 1**). We categorized a total of 48 variants as **ASD** (15 *de novo*) found in cases of ASD, ID, or DD. A set of 4 MS variants were classified as **Somatic Cancer** which have not been reported in ASD, but exhibit a high frequency of reports in the COSMIC database^23^. A further 19 variants termed **PHTS** were found in individuals with PHTS, but not ASD and low frequency in somatic cancer. There is high overlap in variant incidence across these disorders (**Fig. 1b**). We included 5 PTEN variants with well-characterized disruptions on protein function, termed **Biochemical Variants**. These include C124S, which is both protein- and lipid-phosphatase catalytically inactive^24^; G129E, a lipid phosphatase-dead variant^25^; Y138L, a protein phosphatase-dead variant^26^; 4A, in which the four phosphorylation sites Ser^380^, Thr^382^, Thr^383^ and Ser^385^ were replaced with alanines, creating a phospho-null construct lacking self-repression, rendering it constitutively active^27^; and C124S-4A which contains both catalytic inactive and phospho-null mutations^22^. C124S has also been found in somatic cancer^23^, and G129E has been found in macrocephaly, DD^28^, PHTS^29^, and somatic cancer^23^. In order to understand the range of PTEN protein function existing in the general population, 8 variants were selected based on their relatively high frequency in the gnomAD database^30^, termed **Population Variants**. Additionally, in order to test the quality of common predictive algorithms of protein dysfunction, variants were included with either **Predicted High Impact** (17) or **Predicted Low Impact** (5) as determined by CADD phred version 1.0 ^31^ or SNAP2 scores^32^. We also included 2 NS mutations, R130X and R335X, due to their common association with ASD, PHTS and somatic cancer. Many variants exist in multiple categories, and the full variants classification, reference, and CADD phred and SNAP2 scores can be found in **Supp. Table 1**. We use the NCBI reference sequence of PTEN (NM_000314.7) as wildtype (WT).

**Fig. 1.**
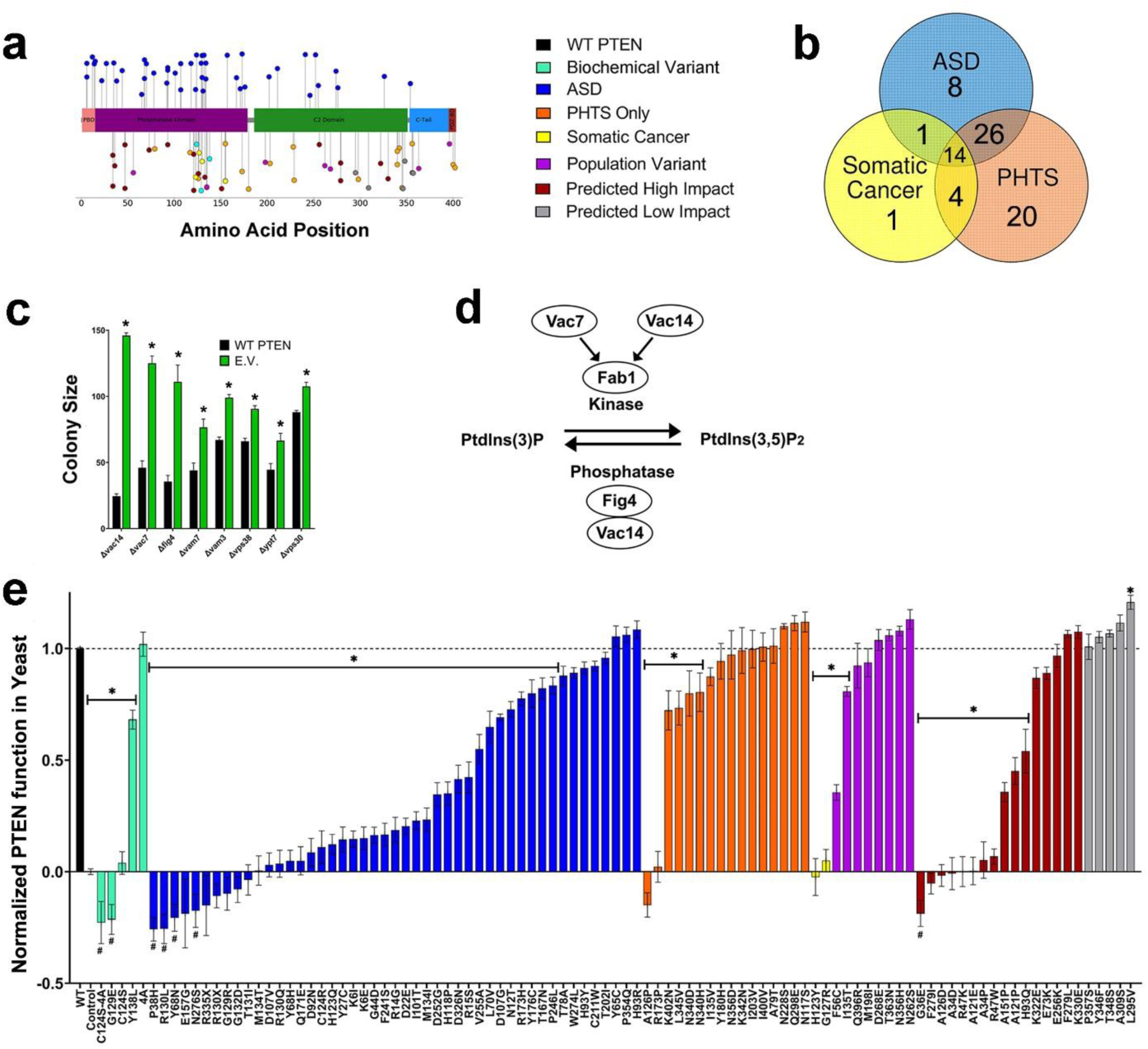
Functional assessment of *PTEN* variants for genetic interactions in yeast. **a** Schematic of 106 MS and NS variants indicating their positions across the functional domains of PTEN. **b** Venn diagram showing the overlap of PTEN variants used in this study identified in individuals with different disorders. The ASD category includes DD; and Somatic Cancer includes variants with 8 or more reports in COSMIC. **c** Overexpressing WT-*PTEN* in a library of 4699 yeast strains, each lacking one nonessential gene, identified 8 sentinel strains showing significant genetic interaction (5 of which function in phospholipid metabolism; 3 in vacuole fusion). **d.** Schematic of sentinel functions in PIP3 metabolism. **e** 99 *PTEN* variants were expressed in the sentinel yeast strain Δ*vac14* and assayed for colony size. Data are expressed as means ±SEM. * indicates p<0.05 for the contrast between WT and variant in the mixed-effects model (see Methods). EV = empty vector.

### PTEN overexpression in yeast reveals novel genetic interactions and provides assessment of lipid phosphatase function

We first took advantage of the high-throughput capacity of the *Saccharomyces cerevisiae* synthetic dosage lethality screen^33^ as an unbiased assay to identify genetic interactions of PTEN. By overexpressing human WT PTEN (WT), empty vector (EV), or the catalytically-inactive variant C124S, in approximately 5000 non-essential gene mutant strains of the yeast deletion collection^34^, we identified 44 potential genetic interactions of PTEN. Through further testing, 8 gene deletion mutants, deemed ‘*sentinels’*, were selected for inclusion in our assay which show significant, strong negative genetic interactions with WT, but not EV or C124S (**Fig. 1c**). In each of these deletion strains, overexpression of WT resulted in a reduction in colony growth. The sentinel proteins Vac7 and Vac14 physically interact and promote the synthesis of PI(3,5)P2 by activating Fab1, a kinase for PI3P^35,36^ (**Fig. 1d**). The sentinel protein Fig4, when recruited to the vacuolar membrane by Vac14, dephosphorylates PI(3,5)P2 to form PI3P^36^. Therefore, lack of any of these three genes results in altered PI3P metabolism. Likewise, the other 6 deletion mutant proteins directly or indirectly influence PI3P signaling^37^. For example, Vam3 and Vam7 are SNARE proteins and Ypt7 is a Rab GTPase. All three are components of the vacuolar fusion complex^38–41^, in which Vam7 directly binds PI3P on the vacuolar membrane to initiate fusion^42^. While yeast does not contain the primary PTEN substrate, PIP3, nor an endogenous PTEN homolog, PTEN also binds PI3P, which is present in yeast, via its C2-domain in the recruitment of cytoplasmic PTEN to the endosomal membrane^43^. Altogether, we hypothesize that PTEN utilizes PI3P as a substrate in yeast and antagonizes already compromised PI3P levels in these mutants.

Next, we designed a novel mini-array platform to screen the genetic interactions of WT, EV, and 99 PTEN variants with each of the 8 sentinel strains. As a representative data set, we present results for the yeast strain lacking Vac14 (*Δvac14*), which shows the strongest genetic interaction with PTEN and allows for the greatest dynamic range to study the effects of PTEN variants (**Fig. 1e; t**hese and all other results also listed in **Supp. Table 2**). We find that using the *Δvac14* sentinel, 37 variants, including 22 of 47 ASD-associated variants tested, exhibit complete LoF, with 6 being below LoF. 27 variants retained partial function while 35 variants, including 8 **ASD**, retained WT-like functionality. Similar results were found with the 7 other sentinels (**Supp. Fig. 1; Supp. Table 3**).

### PTEN variants delay developmental rate in Drosophila

We generated 88 transgenic lines of *Drosophila melanogaster* expressing WT human PTEN, 86 PTEN variants, and an EV control, each integrated into the *attP2* locus^44^ allowing confident quantitative comparison of relative function between all variants by virtue of their equivalent expression levels. In *Drosophila*, reduced insulin receptor signaling slows developmental rate, causing delay in pupa formation and eclosion - the emergence of the adult fly from the pupa^18,45^ (**Fig. 2a**). Given PTEN’s function in counteracting insulin signaling, we found that overexpression of WT in *Drosophila* in all cells throughout development, led to a significant delay in time to eclosion compared to EV, which was not observed in flies expressing C124S (**Fig. 2b**). We then screened 86 PTEN variants, including 45 **ASD** to compare time to eclosion against one another, EV (normalized as 0) and to WT (normalized as 1) (**Fig. 2c**). Variants exhibited a range of functionality: with 41/86 variants being complete LoF, including 26 **ASD**. 34/86 were partial LoF, or hypomorphic, and 11/86 variants retained WT functionality. We also observed 2 variants, the **PHTS** N117S and Q298E, which slowed developmental rate further than that of WT, while the known gain of function (GoF) variant 4A induced lethality. Finally, we noted 6/86 variants, including the lipid phosphatase dead G129E exhibited eclosion significantly faster than EV. We find a high level of concordance (Pearson r = 0.69, p<0.0001) between functional measures of *Δvac14* genetic interactions in yeast and development in fly (**Fig. 2d**).

**Fig. 2.**
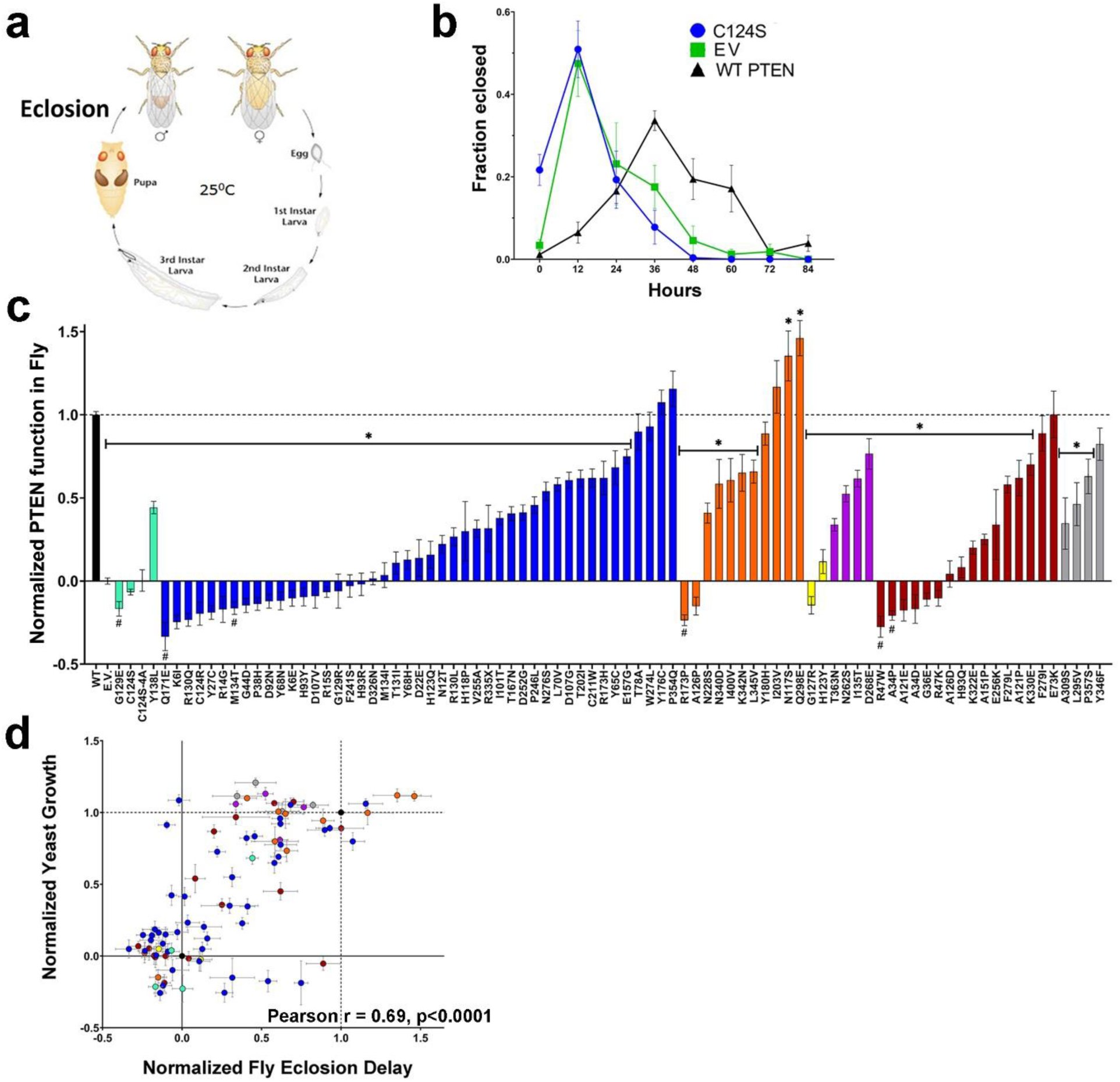
Functional assessment of *PTEN* variants in whole organism development in *Drosophila*. **a,b** Transgenic *Drosophila* overexpressing WT-*PTEN* exhibited delayed development, indicated by increased time to reach eclosion, the emergence of the adult from the pupal sack. **c** Effect on time to eclosion for 88 strains of transgenic flies each expressing a different *PTEN* variant. Data are expressed as means ±SEM. * indicates p<0.05 for the contrast between WT and variant in the mixed-effects model (see Methods). EV = empty vector. **d** Correlation of genetic interaction values from yeast sentinel Δ*vac14* and developmental delay in *Drosophila* for 84 PTEN variants. (Pearson r = 0.69, p<0.0001).

### PTEN variants impact neural development in rat primary cultures

Aberrant neuronal morphology and excitatory/inhibitory synapse balance are hallmarks of ASD^46^, and ASD rodent models^20^. Reducing *PTEN* expression results in increased neuronal growth, with larger soma size, increased dendritic and axonal branch length, and increased excitatory, but not inhibitory synapse density^22,47–51^. To assay variant function in these ASD-associated neuronal growth processes, we overexpressed WT and 19 PTEN variants, or GFP alone, in rat primary hippocampal and dorsal root ganglion (DRG) neuronal cultures. We find that overexpression of WT induced the opposite effects observed by reducing PTEN, including decreased density of PSD95 puncta, a marker of excitatory glutamatergic synapses (**Fig. 3a,c**), but had no impact on density of the inhibitory GABAergic synapse marker Gephyrin (**Fig. 3a**). WT induced a significant reduction of soma size and both total dendritic arbor and axonal lengths (**Fig. 3b,d-f**). For PSD95 density, 14/19 variants including C124S showed complete LoF indistinguishable from controls only expressing GFP, and only 4/19 variants retained WT-like function. In addition, 11/19 variants, exhibited complete LoF activity on dendrite growth, while 7 retained WT-like function. For axonogenesis, 6/19 variants were complete LoF while 13/19 were WT-like. For soma size, variants displayed a varying amount of dysfunction, with 10 variants exhibiting significant LoF compared to WT. C124S and two ASD variants, I101T and G132D, increased soma size more than GFP. The **Population Variant** D268E appeared WT-like in all four measures of neuronal growth and synaptogenesis. Notably, the **PHTS** variant A79T, which is also found at increased frequency in gnomAD, exhibited a GoF phenotype in axonal growth and complete or less than LoF for PSD95 density and dendrite length.

**Fig. 3.**
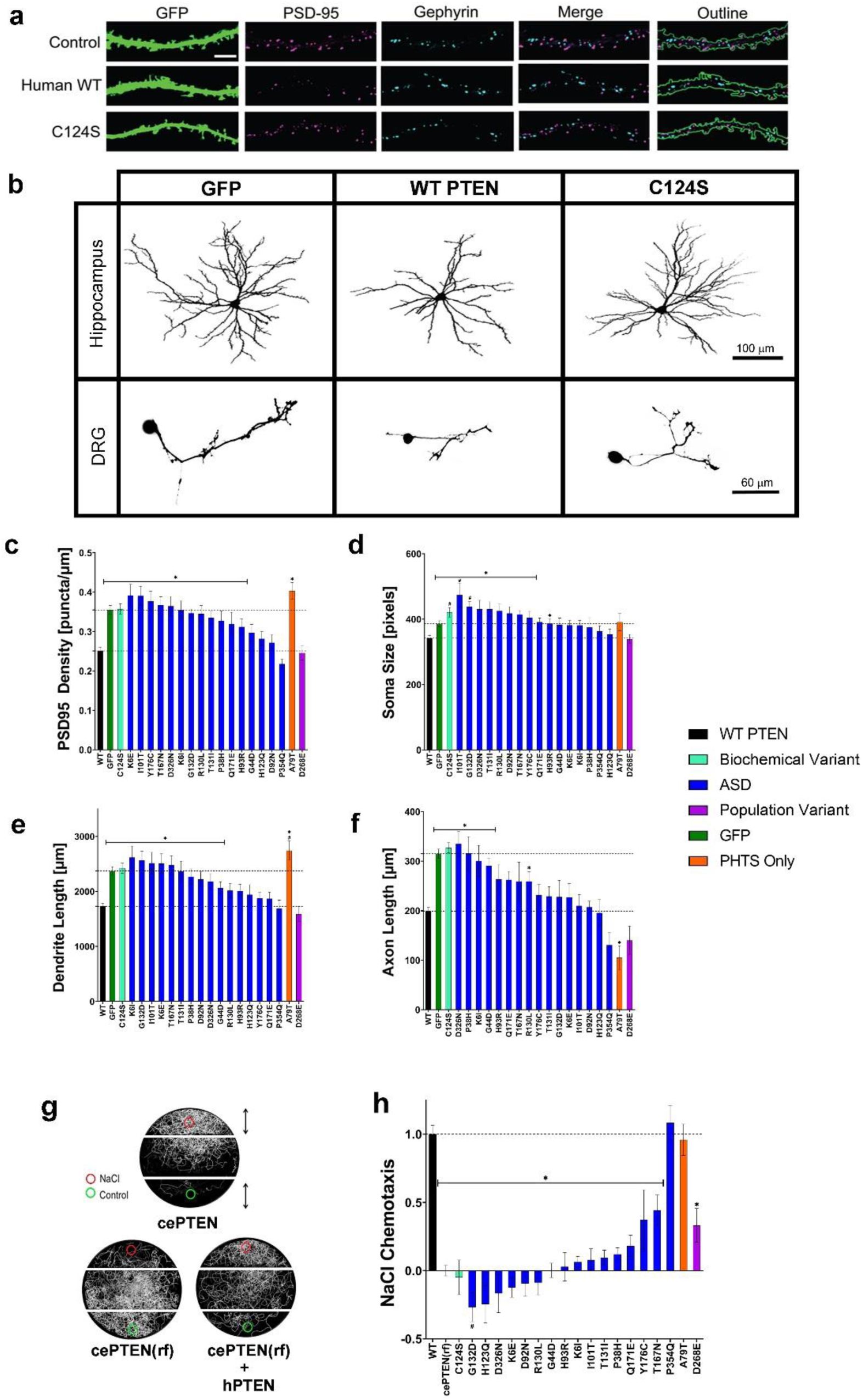
Functional assessment of 20 *PTEN* variants impact synaptogenesis and morphogenesis in rat hippocampal and dorsal root ganglion (DRG) neuronal cultures and sensorimotor behavior in *C. elegans*. **a** Immunofluorescence images of proximal dendrites from primary cultured hippocampal pyramidal neurons co-stained for the postsynaptic markers of excitatory (PSD-95) and inhibitory (Gephyrin) synapses. **b** Representative images of cultured hippocampal and DRG neurons expressing GFP alone, or with WT-*PTEN* and *PTEN*-C124S. Effect of overexpression of 20 *PTEN* variants on excitatory synapses (**c**), soma size (**d**), total dendritic arbor length (**e**) in hippocampal neurons, and axonal length in DRG neurons (**f**). Data are expressed as means ±SEM. * indicates p<0.05 as in **Fig. 1**. **g** Wild type *C. elegans* (cePTEN) exhibit chemotaxis towards a NaCl source, while worms harboring a NS mutation in the worm homolog *daf18* (cePTEN(rf)) exhibit chemotaxis deficits, which is rescued by overexpression of human PTEN (cePTEN(rf)+hPTEN). **h** Impact of 20 *PTEN* variants on NaCl chemotaxis in *C. elegans* is shown by scoring salt preference as (A-B)/(A+B). Data are expressed as means ±SEM and are normalized to cePTEN(rf)+hPTEN=1 and cePTEN(rf)=0. * indicates p<0.05 compared to WT, # marks p<0.05 compared to cePTEN(rf).

### PTEN variant effects on sensory processing and behavior in C. elegans

Since ASD is a behaviorally diagnosed disorder, and because deficits in sensory processing are a core feature of ASD present in >95% of cases^58^, we tested the effects of PTEN variants on sensorimotor neural circuit function in an intact animal model. We used a behavioral assay in *C. elegans* as they have a unique advantage of being the only animal known to survive homozygous null mutations in PTEN, allowing us to directly assess the functional consequences of PTEN variants in a transgenic rescue assay *in vivo*^52^. *C. elegans* exhibit robust navigation up a concentration gradient toward a point source of NaCl. Worms harboring NS mutations in the *PTEN* homolog, *daf-18*, exhibit a complete reversal in NaCl chemotaxis due to a dysregulation in the insulin pathway acting on a single sensory neuron (**Fig 3g**)^59,60^. Pan-neuronal expression of WT human PTEN in *daf-18* mutants was sufficient to rescue normal chemotaxis. We then used our machine vision system, the Multi-Worm Tracker^53^ to automatically quantify chemotaxis in large populations of animals. We overexpressed the same WT and 19 PTEN variants tested in rat neuronal cultures in *daf-18* mutant worms and found that 14/19 variants, including C124S exhibit complete LoF, while T167N, Y176C and D268E showed partial LoF, and A79T and P354Q retained WT-like function (**Fig 3h**). The ASD variant G132D exhibited stronger negative chemotaxis than *daf-18* mutants.

### Protein instability is a major mechanism of variant PTEN dysfunction

To identify molecular mechanisms underlying variable effects of MS variants, we measured impact on protein stability, a known mechanism of PTEN dysfunction^15,27^. We first measured PTEN protein abundance for WT and 97 variants tested in yeast using western blot analysis (**Fig. 4a**) and found a significant reduction in PTEN levels for 31 variants compared to WT, with 21 variants having stability below 50% of WT. Notably, of 46 **ASD** variants measured, nearly half (22/46) had significantly reduced levels of PTEN. To examine whether the changes in protein abundance assessed in yeast is reflected in a human cellular environment, we overexpressed WT and 105 variants as sfGFP fusion proteins in HEK293 cells, with the red fluorophore mTagRFP-T co-expressed from the same plasmid. The green-to-red ratio for each cell was quantified using flow cytometry, with a decrease from WT level indicating reduced protein stability (**Fig. 4b-d**). 77/105 variants were significantly less stable than WT, of which 50 exhibited stability of 50% or less than WT. 19/105 showed WT-like stability, and 9/105 were significantly more stable. Of the 47 **ASD** variants tested, most (40/47) were significantly less stable than WT, and 29 exhibited less than 50% WT stability. Only 5, P354Q, R130L, R15S, R14G and T78A, were not significantly different from WT. The variants H93Y and K6E were hyper-stable. Interestingly, all of the **Biochemical Variants** exhibited significant protein instability. PTEN stability in yeast and HEK293 cells were well correlated (**Fig. 4e**, Pearson r = 0.55, p<0.001) and results indicate that instability of PTEN protein is a major mechanism of single nucleotide variation-induced dysfunction.

**Fig. 4.**
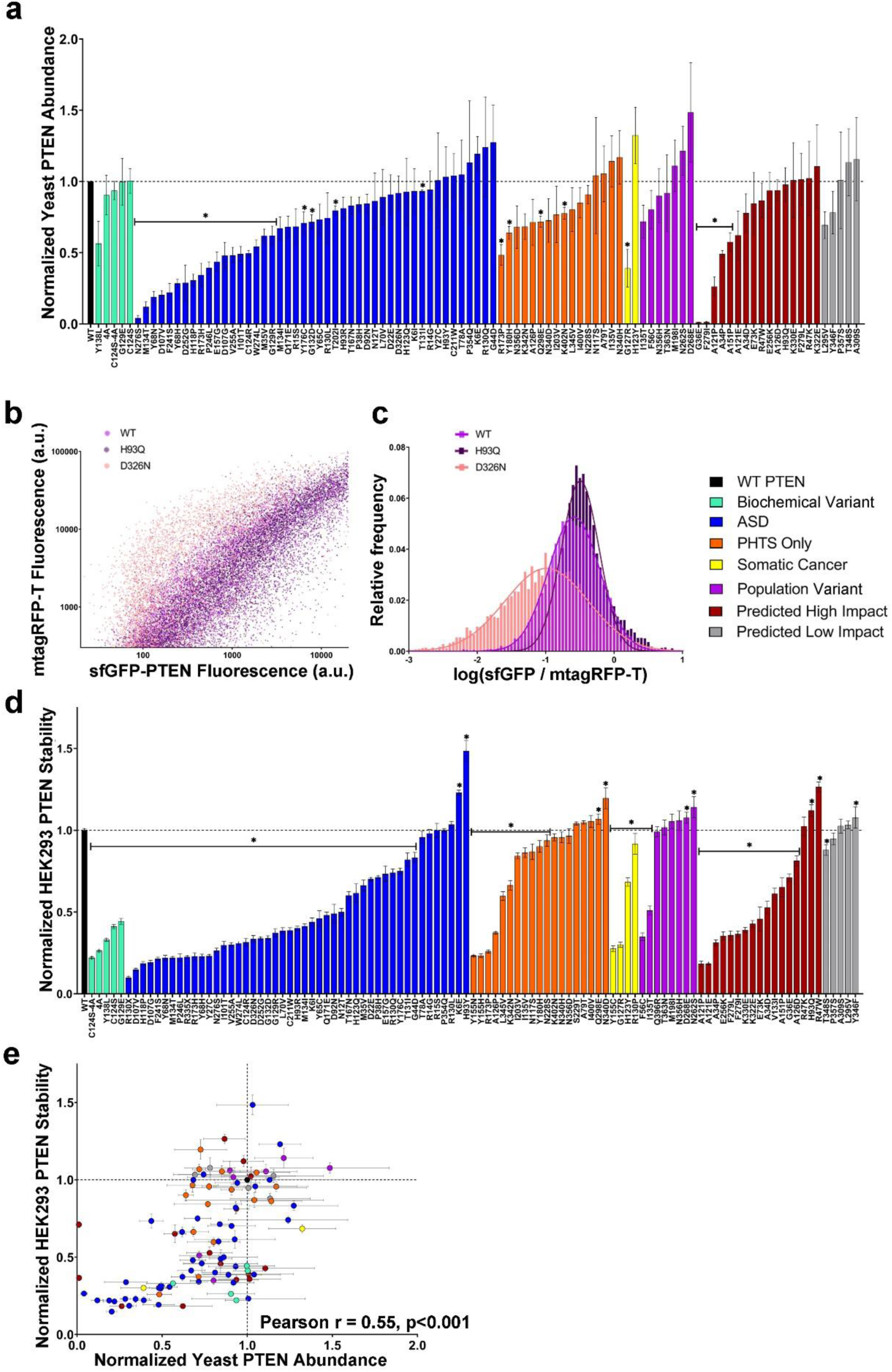
PTEN variants destabilize PTEN. **a** The abundance of 97 PTEN variants overexpressed in yeast was assayed by western blot. Quantified values by band densitometry were normalized to WT-PTEN = 1 and empty vector = 0. **b** Representative scatter plot of single-cell fluorescence intensities from flow cytometry for overexpressed sfGFP-fusions of PTEN WT, D326N and H93Q versus their level of transfection as visualized with mTagRFP-t. **c** Histogram showing the relative frequency of sfGFP/mTagRFP-t ratio for the same variants in (**b**). **d** Protein stability of 105 PTEN variants assayed in HEK293 cells by flow cytometry calculated as median of sfGFP/mTagRFPt and expressed as normalized to WT = 1. Data in **a** and **d** are expressed as mean of well replicates (n ≥ 4) ± SEM. * indicates p < 0.05 compared to WT by student’s t-test. **e** Plot of yeast abundance and HEK293 PTEN stability variant data sets. (Pearson r = 0.55, p< 0.0001).

### PTEN reduction of pAKT/AKT in HEK293 cells identifies 25 novel dominant negative variants

PTEN functions as a negative regulator of the PI3-AKT signaling pathway by decreasing the pool of available PI(3,4,5)P3 via its lipid phosphatase activity, causing a reduction in the level of activated, phosphorylated AKT (pAKT)^54,55^. To assess the lipid phosphatase function of WT PTEN and 105 variants, we measured levels of pAKT/AKT in insulin-treated HEK293 cells using flow cytometry (**Fig. 5a-c**). As expected, overexpression of WT decreases pAKT levels in an exogenous PTEN expression level-dependent manner (**Fig. 5a**). 25/105 variants showed complete LoF and 19/105 showed partial LoF. 29/105 variants retained WT-like function, and P354Q, Q396R and the constitutively active 4A, exhibited a GoF phenotype. Further, 65/105 variants had <50% functionality, including 42/47 **ASD**. Expression of C124S or G129E raised pAKT/AKT levels (**Fig. 5c**), consistent with a dominant negative phenotype, as reported previously^15,21,56,57^. Here, we identify 27 additional variants (25 novel) exhibiting dominant negative activity, including 17 **ASD**. To further investigate these phenotypes, we used CRISPR/Cas9 to create a bi-allelic knockout of PTEN (PTEN-KO) in HEK293 cells and repeated the pAKT/AKT assay for 32 variants. For 21 of 28 variants exhibiting dominant negative activity in parental cells, the absence of endogenous PTEN abrogated this effect (**Fig. 5d**). Interestingly, the GoF of constitutively active 4A was markedly reduced in PTEN-KO, yet still significantly more active than WT, suggesting it may enhance endogenous PTEN activity when present. We found only 6/32 variants tested (A126P, H123Q, P38H, R130Q, A126D and R130L) exhibiting greater than 10% change in stability between parental and PTEN-KO cells suggesting that interactions with endogenous PTEN had minimal influence on variant stability (**Supp. Fig. 2**).

**Fig. 5.**
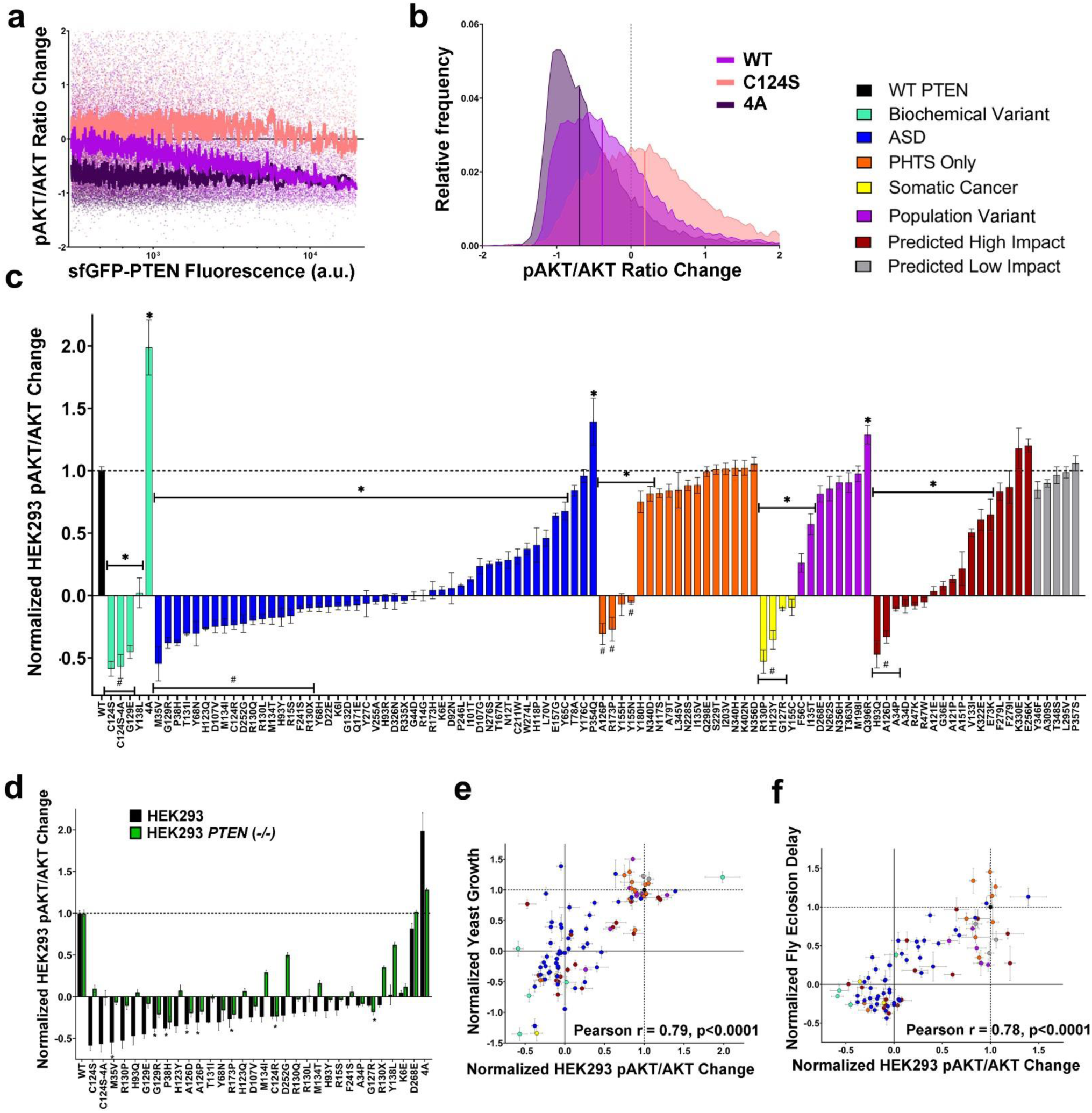
PTEN variant impact on levels of pAKT/AKT reveals LoF and dominant negative activity. **a, b** Flow cytometry single-cell rolling median of pAKT/AKT ratios versus expression levels of sfGFP-PTEN in HEK293 cells for WT, C124S and 4A variants and the corresponding frequency histogram in b). Vertical bars indicate median values. **c** Functional impact of 105 PTEN MS variants on pAKT/AKT levels. Data plotted are median differences between transfected and in-well untransfected cells for each variant Data are expressed as mean of well replicates (n >=4) ±SEM. Values are normalized to WT=1 and no difference to untransfected=0. * indicates p<0.05 compared to WT, # indicates p<0.05 compared to 0 by one-sample Student’s t-test. **d** Relative pAKT/AKT changes are shown for variants exhibiting dominant negative effects on pAKT/AKT, as well as the constitutively active 4A in both parental and a PTEN-KO HEK293 cell line. **e**, **f** Plots of PTEN variant function by pAKT/AKT changes in HEK293 cells versus variant function of genetic interaction with Δ*Vac14* in yeast (Pearson r = 0.79, p< 0.0001), and rate of eclosion in *Drosophila* (Pearson r = 0.78, p< 0.0001).

### Varied functional correlations for variants within stability-dependent or –independent domains

Overall, we find a range of strength of correlation between assays and model systems (**Supp. Fig. 3**), with strongest correlations observed between the 8 yeast sentinels, and between the HEK293 pAKT/AKT assay and yeast genetic interactions (**Fig. 5e;** Pearson r = 0.79, p<0.0001 for *Δvac14*), fly development (**Fig. 5f**; Pearson r = 0.78, p<0.0001), rat axonal outgrowth (Pearson r = 0.73. p<0.001), and worm sensorimotor behavior (Pearson r = 0.82, p<0.0001). Other cross-assay and – model comparisons showed limited concordance.

In order to identify features of variant impact contributing to varied phenotypes in different assays we investigated the relationship between protein function and stability. To determine whether function directly tracks stability we subtracted the normalized stability (nS) of each variant from its normalized function value (nF) in each assay. If function directly follows stability then nF-nS≈0, while values closer to 1 or −1 would suggest function/dysfunction independent of stability. Positive values indicate variants with greater function than predicted from stability, while negative values indicate variants with less function than predicted from stability. We used measures of protein abundance in yeast for nS for yeast, and stability measures in HEK293 for all other assays. Interestingly, we find domains within PTEN in which variants predominantly exhibited either stability-independent function (nF-nS<>0), or stability-dependent function (nF-nS≈0) (**Fig. 6a,b**). Stability-independent domains, encompassing variants which largely exhibit greater dysfunction than predicted from their instability (nF-nS≈-1), are associated with previously well-characterized substrate binding and catalytic domains including within the N-terminal phosphatase and the PIP3-binding domain (PBD) (amino acid (AA) positions 1–55); the WPD-loop (AA 92–93); and the P-loop (AA 123–131)^58^. Therefore, we define stability-independent domains as AA 1–55, 92-93, and 123-131, and variants outside these regions, largely exhibiting nF-nS ≈ 0, as being in stability-dependent domains (**Fig. 6a,b**). Variants within these 2 groups showed distinct distributions when plotting the frequency of nF-nS values (**Fig. 6c**). Plotting the HEK293 pAKT/AKT function against stability reveals that for variants in stability-dependent domains function levels track with stability, yielding a range of function from WT to partial and full LoF (**Fig. 6d**). For these variants, instability is likely the primary molecular mechanism underlying dysfunction. In contrast, variants within stability-independent domains exhibit complete LoF or dominant negativity irrespective of level of instability, likely indicating molecular mechanisms of dysfunction associated with aberrant substrate binding or catalytic activity. We find similar distributions of nF-nS values using data from yeast or combining results from all models (**Supp. Fig. 4**), and when removing variants with function and stability values not significantly different from WT (**Supp. Fig. 4f**).

**Fig. 6.**
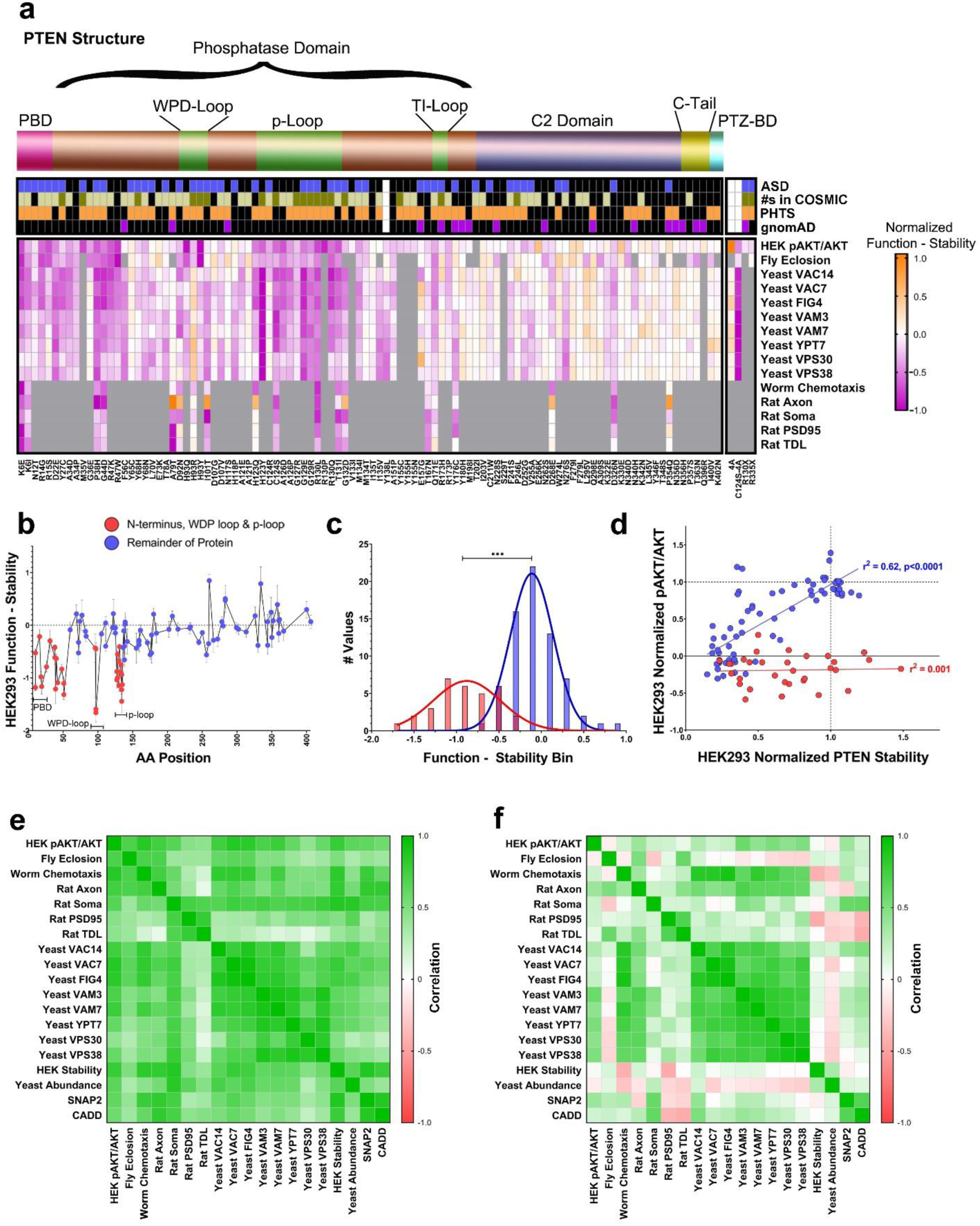
Comparison of functional and stability data identifies instability and loss of catalytic function as distinct mechanisms of molecular dysfunction. **a.** A normalized plot displaying stability values subtracted from function values for each assay with variants displayed according to their amino acid position below a schematic of PTEN structural domains. Normalized function – normalized stability scores are depicted as a heat map in which a score of 0 (white) indicates variants whose function matches their stability. A positive score (orange) indicates higher function than predicted from instability, while negative scores (magenta) indicates greater dysfunctional than instability. **b** Variant function-stability scores for pAKT/AKT assay in HEK293 cells plotted against amino acid position with variants separated into N-terminus, WPD-loop and P-loop domains (red), and variants outside these domains (blue). **c** Frequency distribution of normalized function - stability scores for pAKT/AKT assay in HEK293 assay showing two distinct populations, with red and blue bars denote variants as in b). **d** HEK293 pAKT/AKT assay data plotting PTEN variant function vs. stability with data separated by variants as in b): functional dysfunction of variantsthat is explained by stability (blue), and not (red). **e** High correlations are found for variant impact on multiple assays for variants in which dysfunction is associated with instability (blue variants in **b**, **c** and **d**). **f**. Weaker cross-assay correlations are seen for variants in catalytic domains which exhibit greater dysfunction then explained by instability (red variants in **b**, **c** and **d**).

Repeating correlation analyses for variant impact between assays produces dramatically different results when comparing variants within stability-dependent and -independent domains (**Fig. 6e,f; Supp. Fig. 5a,b**). Correlations were high between all assays for variants within stability-dependent domains, indicating that mechanisms of protein instability are independent of model system. Correlations for variants in stability-independent domains, however, were poor across models, suggesting that different assay are selectively sensitive to dysfunction of different substrate binding and catalytic domains.

### Relationship of PTEN variant dysfunction and disease

Attempts to associate MS induced dysfunction to specific disease states is a major role for functional variomics for PTEN, however, this goal is challenged by variants selection bias and high overlap of the same variants occurring in different disorders. To investigate this relationship with data from our multiple assays, we reclassify variants as **ASD** or **PHTS** if they have been identified in any individual with these disorders (including ID or DD in **ASD**), and as **Somatic Cancer** if variants have 8 or more cases reported in COSMIC. For this analysis, we allow the same variants to be in multiple classes. Of the 106 PTEN variants we tested, 49 have been identified in individuals with ASD, DD or ID, with 40/49 also found in PHTS, 15/49 in somatic cancer, and 14/49 in all three disorders (**Fig. 1b**; **Supp. Table 1**). Further, 64 of the total 106 are **PHTS**, 20 are **Somatic Cancer**, and 18 found in both disorders. We find that in yeast and HEK293 lipid phosphatase assays, **ASD**, **PHTS** and **Somatic Cancer** variants all performed significantly worse than **Population Variants.** However, no significant difference was found for variant function between these disorders, except between **PHTS** and **Somatic Cancer** in the HEK293 pAKT/AKT assay (**Supp. Fig. 6a**), but this difference was lost when variants we determined to be ‘likely benign’ due to WT stability and function in multiple models (below) were removed from analysis (**Supp. Fig. 6b**). We note that this analysis relies on assumptions of disease causality that are uncertain, so we interpret the negative result as inconclusive on the question of whether there are disease-specific differences in PTEN variant effects. In contrast to between-disorder comparisons, we find that variant impact on function in the yeast, fly and HEK293 assays all show significant predictive powers when stratified by their frequency of occurrence in somatic cancers as reported in COSMIC (**Fig. 7**).

**Fig. 7.**
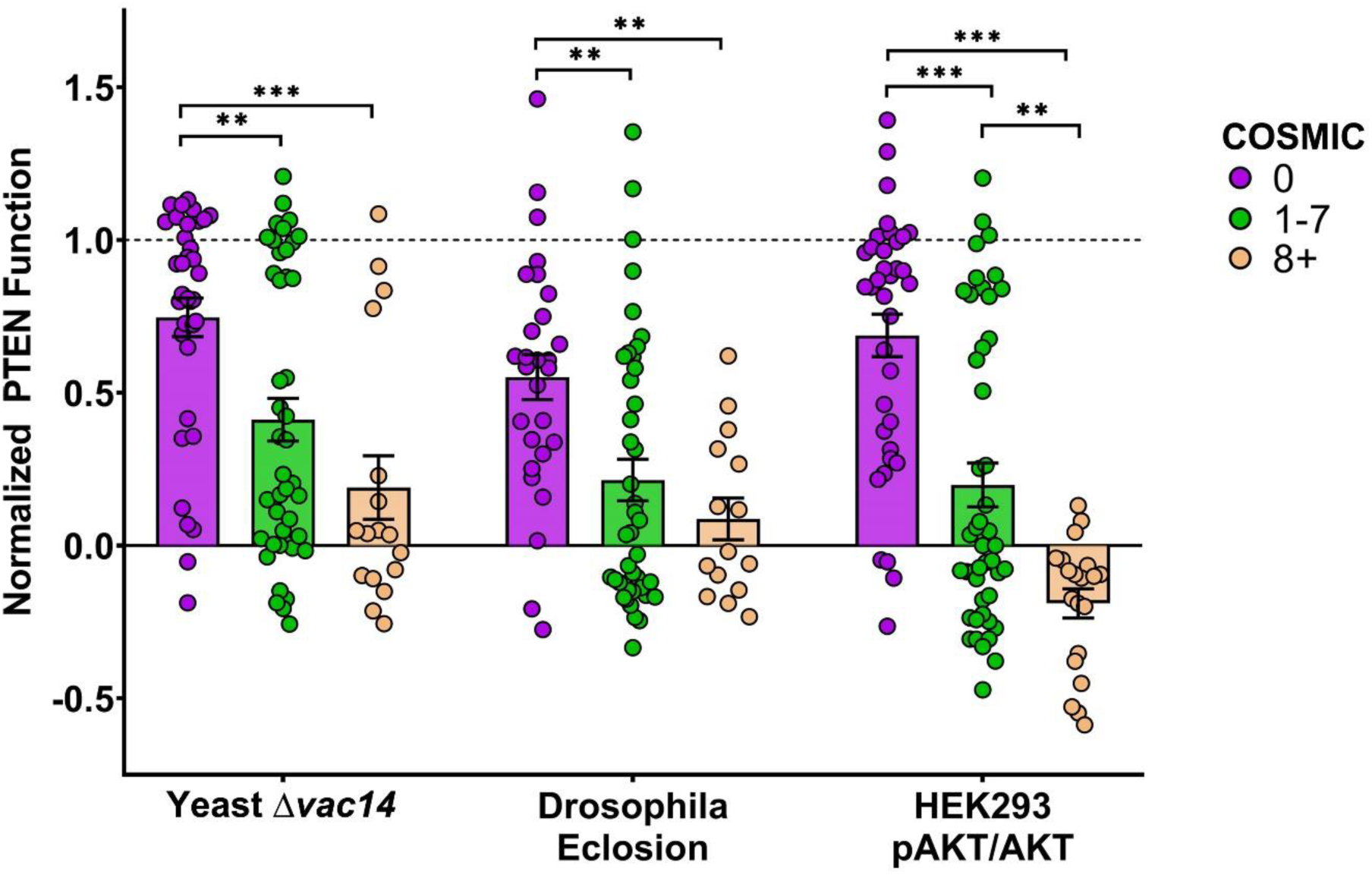
Severity of variant dysfunction correlate with occurrence in cancer. Distribution of normalized PTEN variant function by its frequency within the COSMIC database. In **a** and **b**, mean PTEN function is plotted as bars ± SEM. * p<0.05, ** p<0.005, *** p<0.0005 by two-way ANOVA.

### Assessment of predictive algorithms

We set out to test the value of algorithms for predicting variant dysfunction by testing variants with predicted low or high LoF in CADD phred and SNAP2, computational tools that integrate multiple annotations to quantitatively predict deleterious mutations^31,32^. For variants in the **Predicted Low Impact** group, both CADD and SNAP2 scores were accurate for *Δvac14* yeast, and HEK293 pAKT/AKT assays, but are less accurate predicting results of the fly eclosion assay (**Supp. Table 2; Supp. Fig. 7**). The accuracy of predictions for **Predicted High Impact** variants were more variable. In general, CADD and SNAP2 scores correlate well with our functional assessments, but both are biased toward reporting dysfunction. For example, in ASD variants D107G, L70V, T167N, and Y65C, CADD predicts they are more pathogenic than at least 99% of all other mutations in the genome while SNAP2 predicts a more intermediate effect. However, these variants have WT-like levels of function in our assays.

### High-confidence variant classification

We classified 106 PTEN variants according to their impacts on 8 functional assays encompassing a range of phenotypes between dominant negative to GoF, and 2 abundance/stability assays (**Fig. 8a**). Next, we classified variants following a binary LoF/WT-classifications with a cut-off for LoF at <50% of WT function^59^ (**Fig. 8b**). For each variant, a score from 0 to 1 is then obtained by dividing the number of LoF assays by the number of assays performed, and then classified as follows: ***Likely Benign*** variants have scores <0.25 and are WT-like in at least two assays; ***Likely Pathogenic*** have scores > 0.5 and are LoF in at least two assays; ***Pathogenic*** have scores ≥0.75 and are LoF in at least three assays; and ***VUS*** have scores ≥0.25, but <0.5 or too little assay data yielding conflicting results. Stability was only included in discerning variant profiling for unstable proteins, since WT-like stability is not predictive of WT-like function, but instability correlates with dysfunction.

**Fig. 8:**
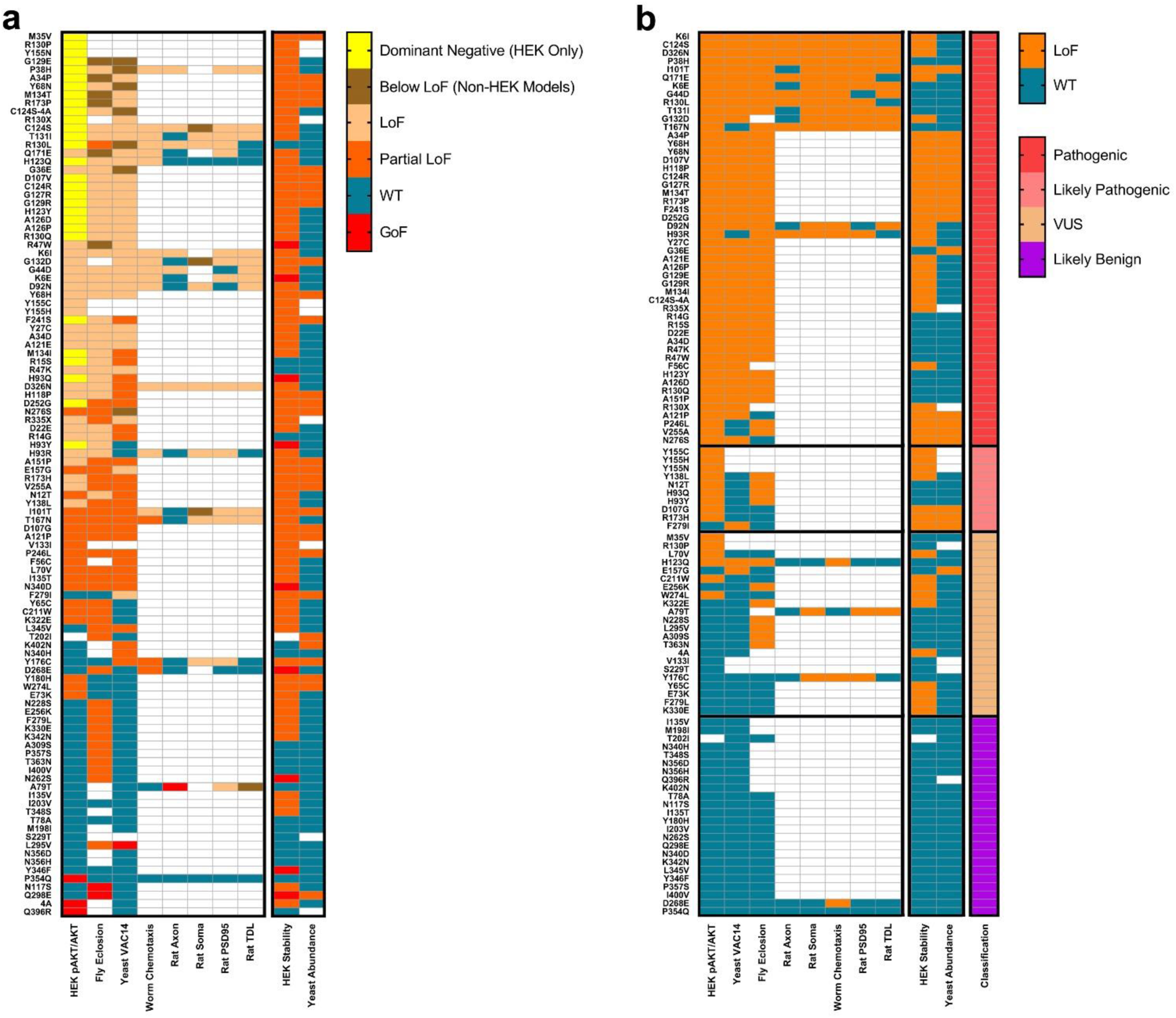
PTEN variant classification based on 9 assays of variant functions. **a** The results of 9 functional assays for 106 PTEN variants are summarized according to the following criteria: WT-like variants exhibit no differences to WT, GoF exhibit function values significantly greater than WT, Partial LoF exhibit function significantly greater than null but significantly less than WT, Complete LoF exhibit no differences to null, dominant negative exhibit function significantly less than both null and WT (Below LoF for fly and yeast assays). Classification are based on statistically significant differences of at least p<0.05 compared to the WT-PTEN, null or both. White cells indicate that the variant was not tested in that assay. **b** PTEN variant predicted impact based on a binary notation of either LoF (<50% of WT effect) or WT-like (≥50% of WT effect), and classified accordingly based on their frequency of LoF (pathologenic)or WT (likely benign) in 9 assays.

Of the 106 variants tested, we classify 50 as **Pathogenic**, including 31 **ASD**. We further classify 10 variants, including 4 **ASD**, as **Likely Pathogenic**. We consider 24 variants to be **Likely Benign**, including 3 **ASD**: P354Q, T202I and T78A; and 12 **PHTS**: I135V, N340H, N356D, K402N, N117S, Y180H, I203V, Q298E, N340D, K342N, L345V and I400V. We are unable to make a functional classification for 22 variants due to either insufficient data or conflicting results between assays and thus retain the classification of **VUS**. These include the **ASD** variants M35V, L70V, H123Q, E157G, C211W, W274L, Y176C and Y65C, as well as the **PHTS** variants A79T, N228S and S229T.

## DISCUSSION

We have taken a deep phenotyping approach to comprehensively assess the impact of single amino acid changes to PTEN function. We use 5 phylogenetically diverse model systems and phenotypes spanning molecular function to behavior in order to measure variant impact on the multiple complex roles of this protein. By taking advantage of the high-throughput nature of yeast, fly, and HEK293 models, we are able to conduct in-depth analyses of ∼100 variants, allowing comparison of effects of overexpression in highly diverse cellular environments. These high-throughput assays allowed selection of a smaller subset of variants exhibiting a range of dysfunction phenotypes to test in lower-throughput neuronal and behavioral assays with more direct relevance to ASD pathophysiology. By utilizing a multi-model system approach including measures of protein stability and function, we identify a diversity of variant impact caused by distinct molecular mechanisms, which would not be apparent from single-model approaches.

As previously reported^15^, we find that PTEN variants commonly decrease protein stability. We find that MS variants induce a full range of instability, which was largely consistent between yeast and HEK293 models. Further, for ∼50% of PTEN variants tested, located throughout the protein structure, there was strong correlation between instability and dysfunction, resulting in a corresponding range of partial to complete LoF. The consistency of this correlation across models suggests conserved mechanisms for MS-mediated PTEN degradation. For these variants, reduced protein abundance is likely the molecular mechanism underlying dysfunction in each assay, and if consistent under physiologic expression in humans, similar levels of haploinsufficiency may contribute to disease expression.

Other variants, including some exhibiting instability, levels of stability did not correlate with protein function, indicating additional molecular mechanisms underlying dysfunction. 16 variants, including 4A, exhibited more function than expected from their stability, suggesting GoF with or without instability. However, most of these variants exhibited greater dysfunction than expected from their instability. These variants largely localized to well-characterized substrate binding and catalytic domains of PTEN, including the PIP3-binding and N-terminal region of the phosphatase domain, and the catalytic pocket encompassesing the WPD- and P-loops. Phenotypes were typically complete LoF or dominant negative/greater than LoF, rather than partial LoF, indicating that these functional domains are intolerant to modification.

The large number of variants exhibiting dominant negativity in HEK293 cells was unexpected. While C124S and G129E have previously been established as dominant negatives, such a characterization of other PTEN variants is sparse^15,21,56,57^. Remarkably, we find 29 variants, including 17 **ASD**, showing dominant negativity in measures of activated AKT. Furthermore, P38S and R130G, two different amino acid substitutions at the same positions as MS variants exhibiting dominant negativity here (P38H, R130L, R130P and R130Q), have previously been reported to have a dominant negative effect on pAKT levels^15,56^. We also find 6 dominant negative variants not found in ASD, but identified in PHTS and/or somatic cancer. Identifying dominant negative variants is of significant clinical value since they may produce outcomes distinct from LoF variants producing haploinsufficiency. Whether phenotypes identified in an overexpression assay are conserved at physiological levels, as shown for C124S and G129E^56,57^, requires further study. Overall, these results suggest multiple mechanisms of MS mutation-induced PTEN dysfunction: instability producing haploinsufficiency, and loss of substrate binding or catalytic function leading to either complete LoF or dominant negativity.

We found varied strength of correlations between functional measures of PTEN variants across assays and model systems overall, but high correlations for variants in which dysfunction is due to instability. This suggests conserved mechanisms of MS-mediated protein degradation resulting in reduced protein affecting all assays similarly. Increased intra-assay and -model variability was observed for variants whose dysfunction was associated with mutations in substrate binding and catalytic domains, likely reflecting selective sensitivity of each assay to distinct functions of PTEN. For example, for all 8 sentinel assays in yeast, variants within the WPD were better tolerated than in other model systems, potentially due to differences in steric requirements for interactions with the substrate PI3P in yeast and the primary substrate PIP3 in other models. We also find that the sentinels *Δvac14*, *Δfig4* and *Δvac7*, which mediate PI3P-to-PI(3,5)P2 interconversion, are particularly sensitive to mutations in the PIP3-binding domain, implicating PTEN-mediated lipid interconversion in these assays. The other 5 sentinels, *Δvam3*, *Δvam7*, *Δvps30*, *Δvps38* and *Δypt7*, are more sensitive to variants located adjacent to the CBR3 loop which mediates PTEN binding to endosomal membrane PI3P^43^, potentially implicating sensitivity to PTEN’s role in endosomal trafficking. The fly eclosion assay, however, exhibited heightened sensitivity to mutations in the TI-loop. Other functional domains exhibit conserved sensitivity across models, including the beginning of the phosphatase domain, bordered by variants D22E to R47K and R47W, the PIP3-domain and the P-loop. These results highlight the strength of a multiple-assay and -model system approach for fully assessing variant impact on multifunctional proteins.

Complex patterns of variants’ effects on our measures of neuronal morphogenesis and sensorimotor behavior suggests PTEN has multiple functions in neurons. This is supported by previous findings that both lipid and protein phosphatase PTEN activity participate in synaptogenesis^22,60^. Here, we find similar patterns of effects on measures of excitatory synaptogenesis and dendritogenesis, indicating common PTEN action. Alternatively, since synapse formation directly affects dendritic growth^61^, their similar patterns of effects may be driven primarily by synaptic changes. Such an interaction would not occur in the axonogenesis assay, since axon growth is measured at stages prior to onset of synaptogenesis. For some variants, results from rat neuronal growth and worm behavior assays differed from yeast and fly assay. For example, H123Q has less impact on the neuronal and behavior assays, while greater impact was seen for a number of **ASD** variants, including D326N, T167N, I101T, as well as the **PHTS** variant A79T.

A critical role of functional variomics is establishing whether VUS encode dysfunctional proteins potentially causal to disease, or protein with WT function and likely benign. Given the overlap of specific variants associated with ASD, PHTS and somatic cancer, limited annotation of patient phenotypes, and our inclusion of variants identified in single cases of ASD and PHTS, we could not attribute distinct molecular mechanisms of PTEN dysfunction to distinct disease states. However, of the 74 variants we tested found in ASD, PHTS, and/or somatic cancer, we find 10 with phenotypes not significantly different from WT, or with >50% WT function in all assays tested. We predict that these variants are likely benign and not contributing to disease. For **ASD**, the three variants, T78A, T202I and P354Q, show WT activity across assays. P354Q is also the third-most common PTEN variant identified in the population (gnomAD), further supporting its classification as benign. The remaining **ASD** variants exhibit dysfunction ranging from partial to complete LoF, and dominant negativity. Furthermore, we find a strong relationship between variant impact on multiple assays and their frequency of detection in cancerous tumors. All of the 19 MS variants we examined with at least 8 reports in somatic tumors showed LoF in all high-throughput models, particularly those located proximal to the lipid phosphatase domain. Altogether, our results strongly support a multi-model system approach for high confidence profiling of genetic variants.

## Supporting information

PTEN variant annotation

PTEN variant assay data

Pearson correlations for all assay data

C elegans strains used

## Author Contributions

The manuscript was written by K.L.P and KH; selection and annotation of PTEN variants was performed by D.B.C. and S.R.; the variant mutation library was created by F.M. and W.M.M.; yeast experiments performed by K.L.P. and B.P.Y.; experiments in *Drosophila* were conducted by P.G.; experiments in rat neuronal cultures were carried out by R.D., M.E., and C.H.; experiments in *C. elegans* were performed by T.A.M; HEK293 experiments were conducted by F.M. and W.M.M.; the HEK293 PTEN-KO cell line was created by A.C. using CRISPR/Cas9; and statistical analyses were performed by M.B.

## Acknowledgement

This work was supported by grant from the Simons Foundation (SFARI award #573845, Grantees: KH, PP, DWA, CJL, SXB, TPO, CHR), and a CIHR Foundation Award (KH).

---

**Supplemental Figure 1.**
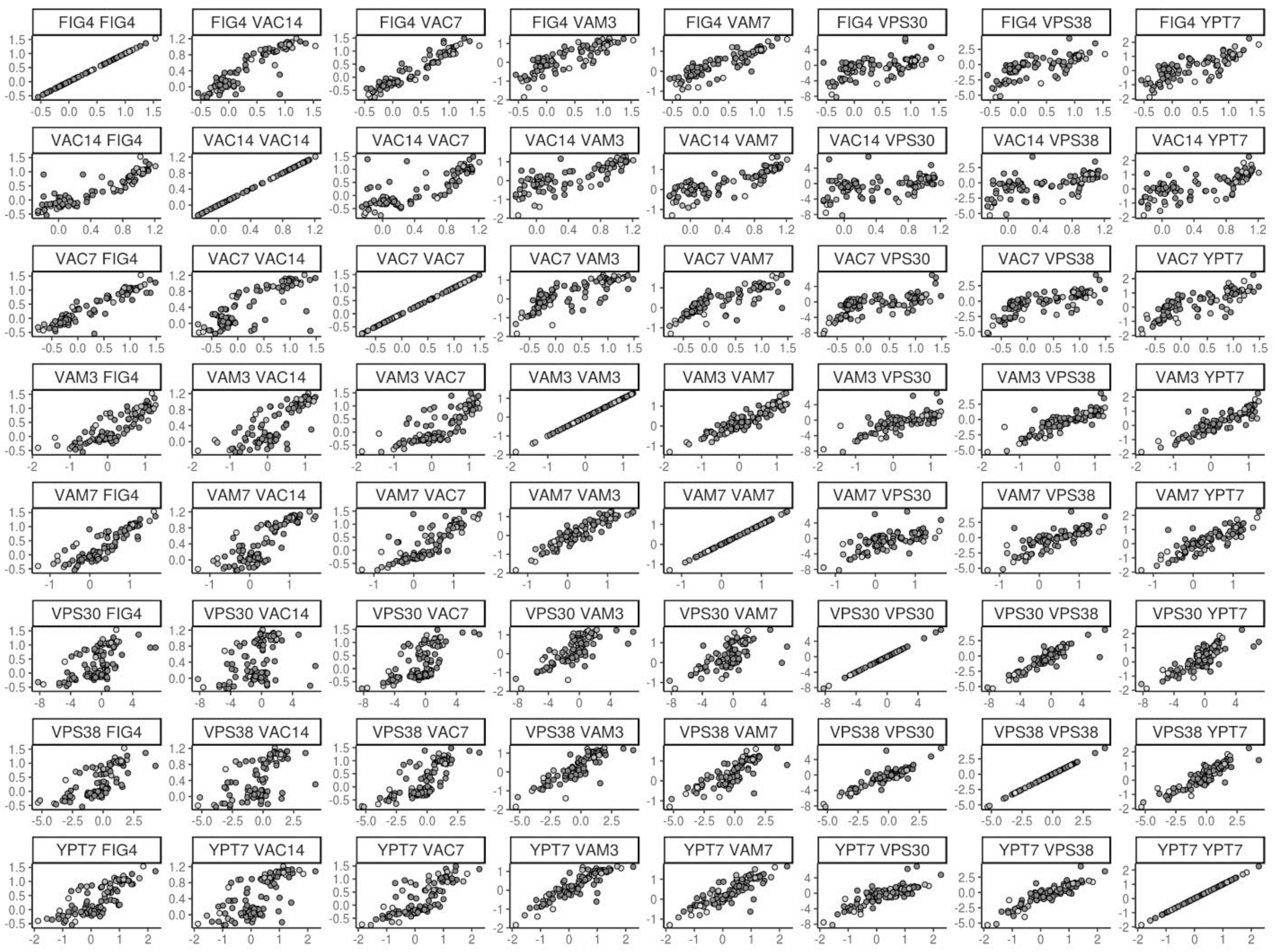
Correlations between 8 sentinel yeast strains. Correlation plot matrix across all eight yeast sentinels assayed for genetic interaction with PTEN. Pearson coefficients and p values in **Supp. Table 3**.

**Supplemental Figure 2.**
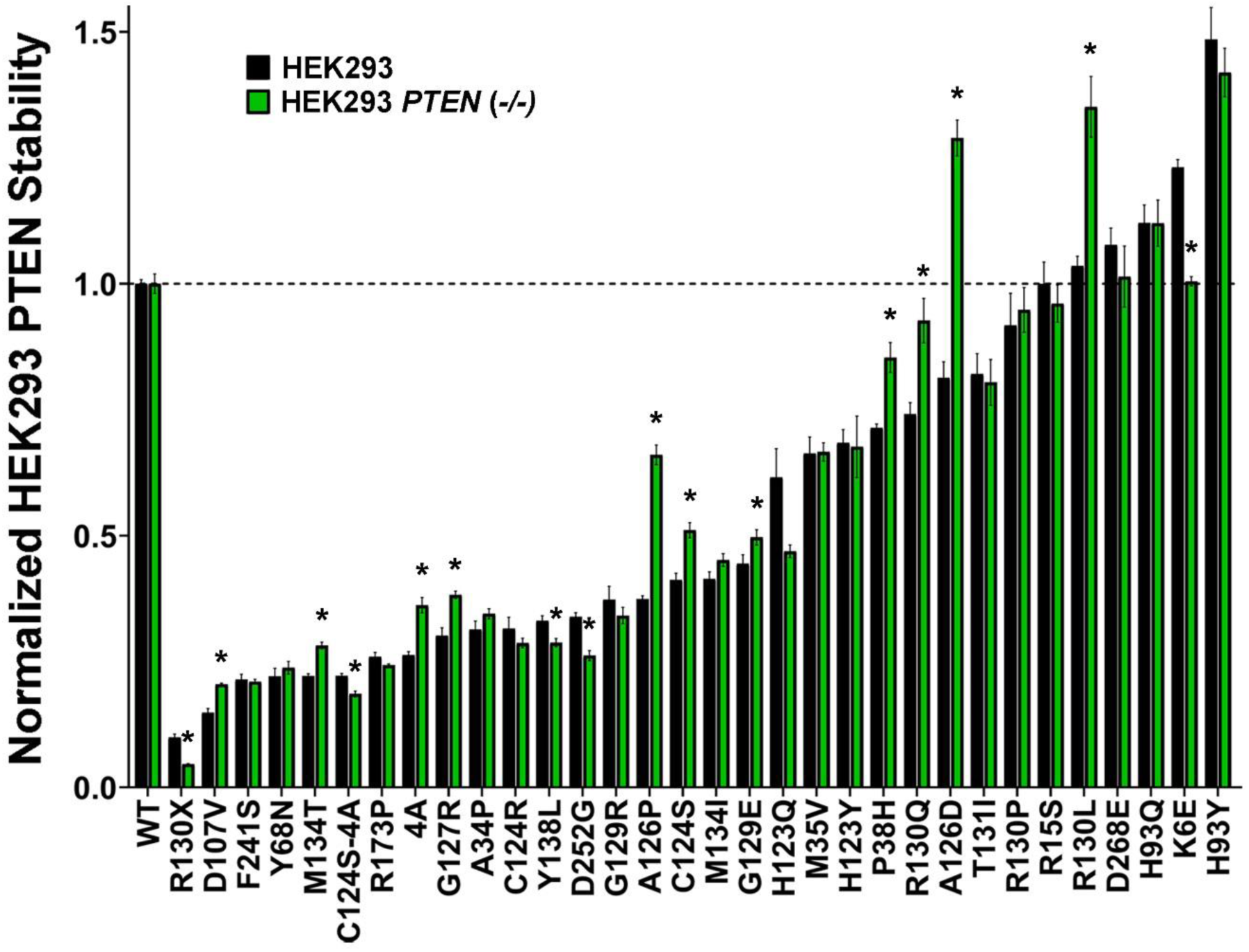
Loss of endogenous PTEN has limited impact exogenous PTEN variant stability. Protein stability of 32 PTEN variants assayed in HEK293 cells and HEK293 PTEN KO cells by flow cytometry calculated as median sfGFP/mTagRFPt and expressed as normalized to WT = 1. Plotted is mean across replicates +-SEM. Asterisk indicates p<0.05 by multiple t-test comparison across cell lines.

**Supplemental Figure 3.**
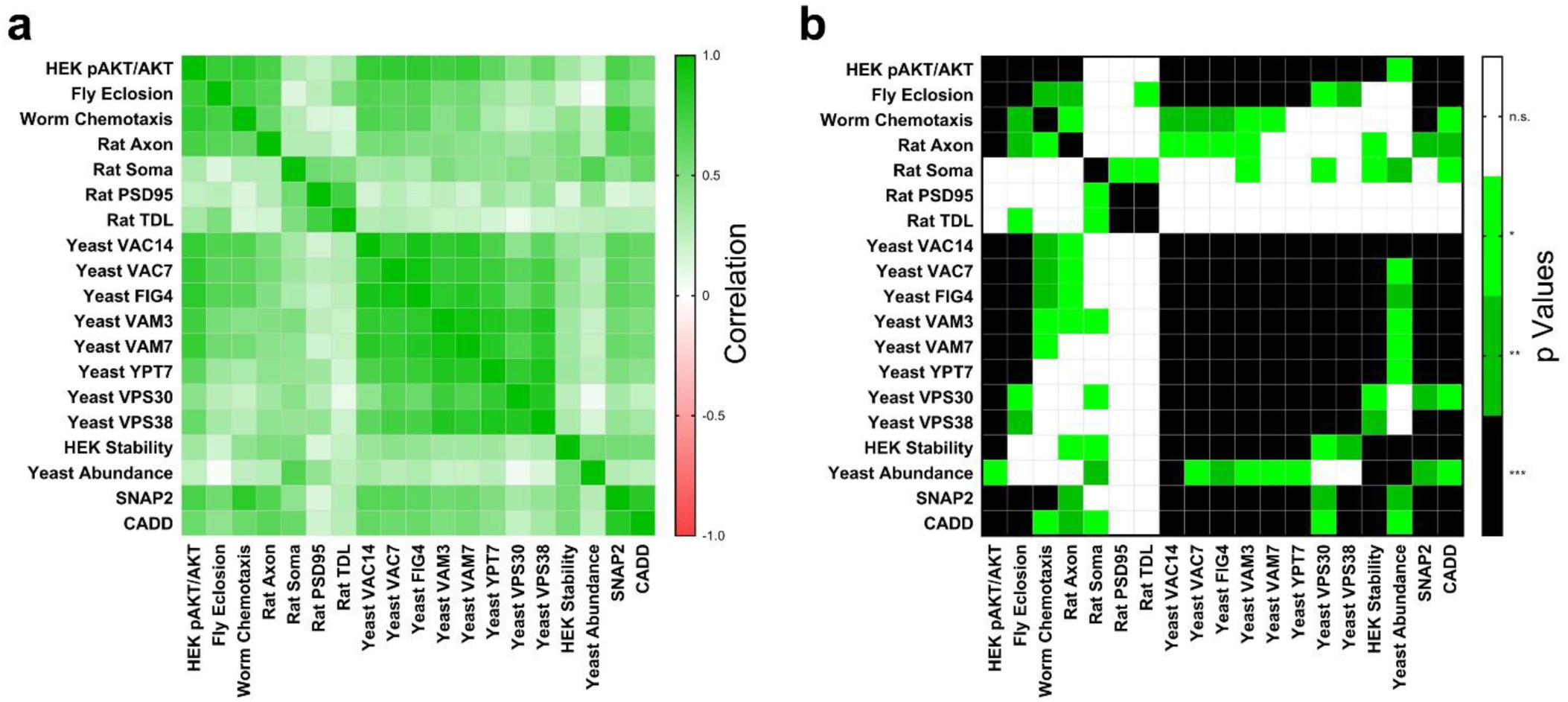
Correlations between 17 assays in 5 model systems. **a** Pearson correlation matrix across model-systems and assays for all variants assayed in each dataset. Strong correlation of the two stability/abundance datasets in yeast and HEK293 with functional metrics across assays. **b** p-value matrix plot of Pearson correlation matrix indicating different significance thresholds.

**Supplemental Figure 4.**
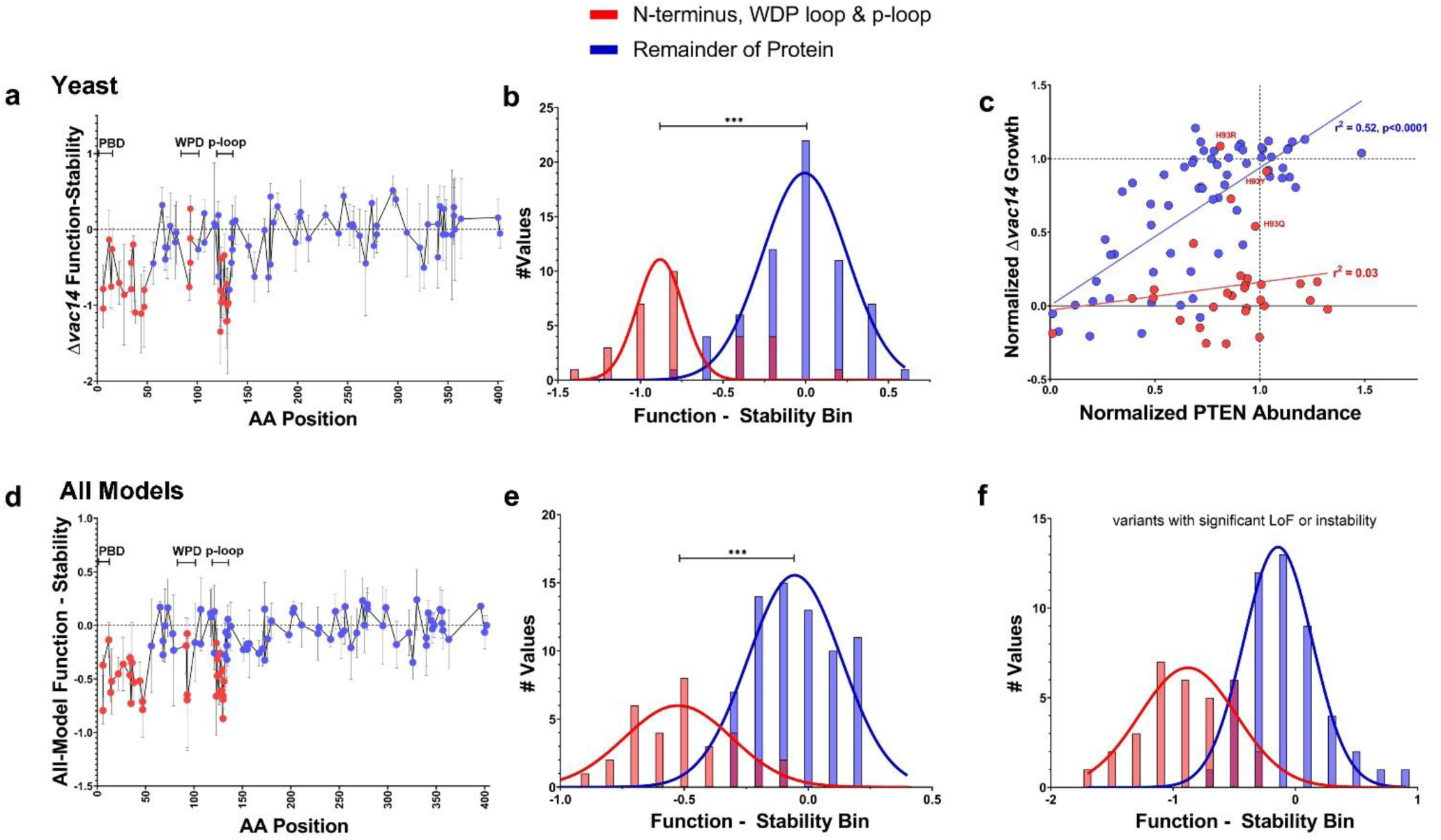
Stability-dependent and –independent PTEN domains in yeast assay and all models. **a** Average variant function-stability (nF – nS) scores for the yeast Δ*vac14* assay plotted against amino acid (AA) position, showing clear separation of variants with low scores (stability-independent) predominantly in the well-characterized domains within the N-terminus, WPD-loop and P-loop (red), and variants exhibiting nF – nS ≈ 0 outside these domains (blue). Abundance in yeast used for nS. **b** Frequency distribution of nF – nS for the yeast Δ*vac14* assay showing two distinct distributions for variants in stability-independent (red) and –dependent (blue) domains. **c** Yeast Δ*vac14* assay data plotting PTEN variant function vs. abundance with data separated by variants within stability-independent (red) and –dependent (blue). **d** Average variant nF – nS scores (using yeast abundance for nS for yeast, and HEK293 stability for all other models) across all assays plotted against AA position with variants separated into variants in stability-independent (red) and –dependent (blue) domains. **e** Frequency distribution of nF – nS for all assays showing two distinct distributions for variants in stability-independent (red) and –dependent (blue) domains. **f** Frequency distribution of nF – nS for all assays as in (**e**), but with removing variants exhibiting WT-like function or stability.

**Supplemental Figure 5.**
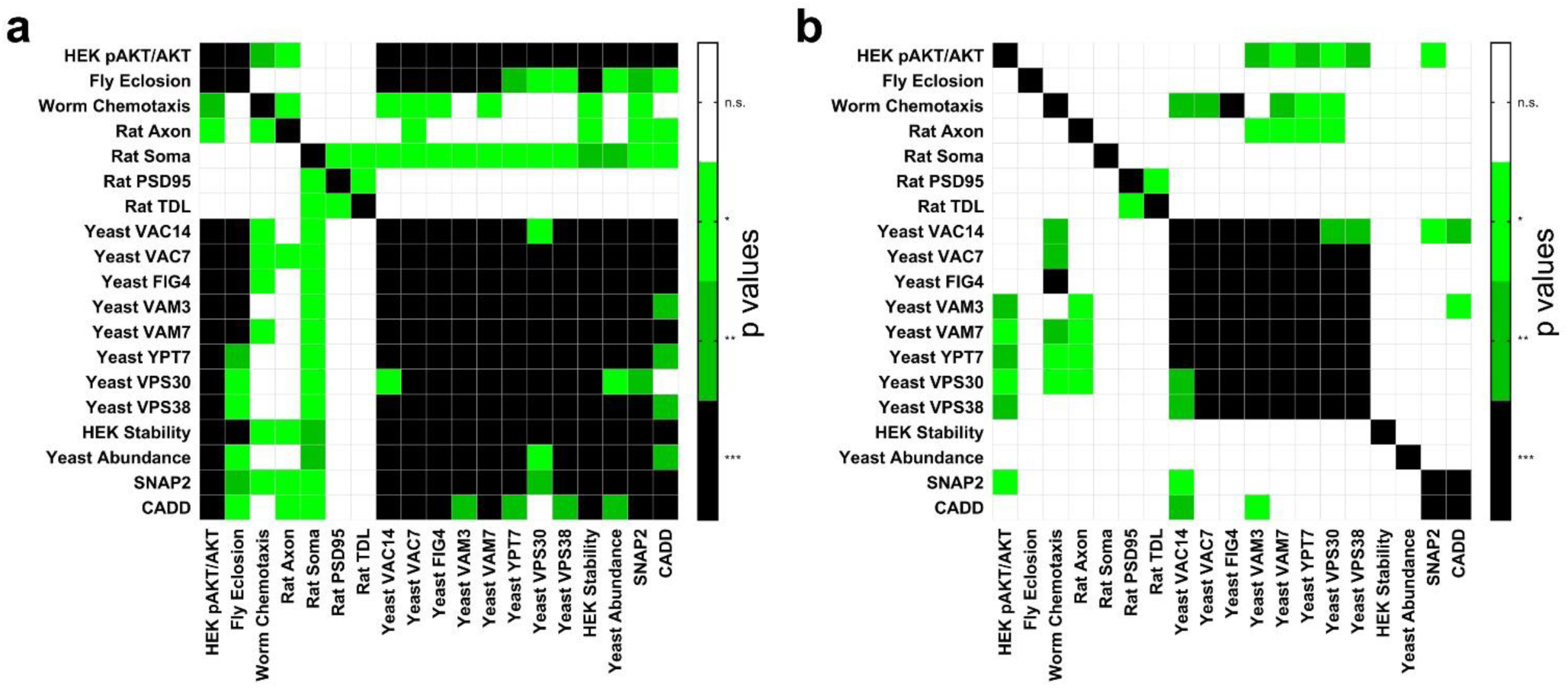
**a,b** p-value matrix plots of Pearson correlation matrices shown in **Fig.s 6e,f** indicating stronger significance between assays for variants in stability-dependent (**a**) compared to –independent (**b**) domains.

**Supplemental Figure 6.**
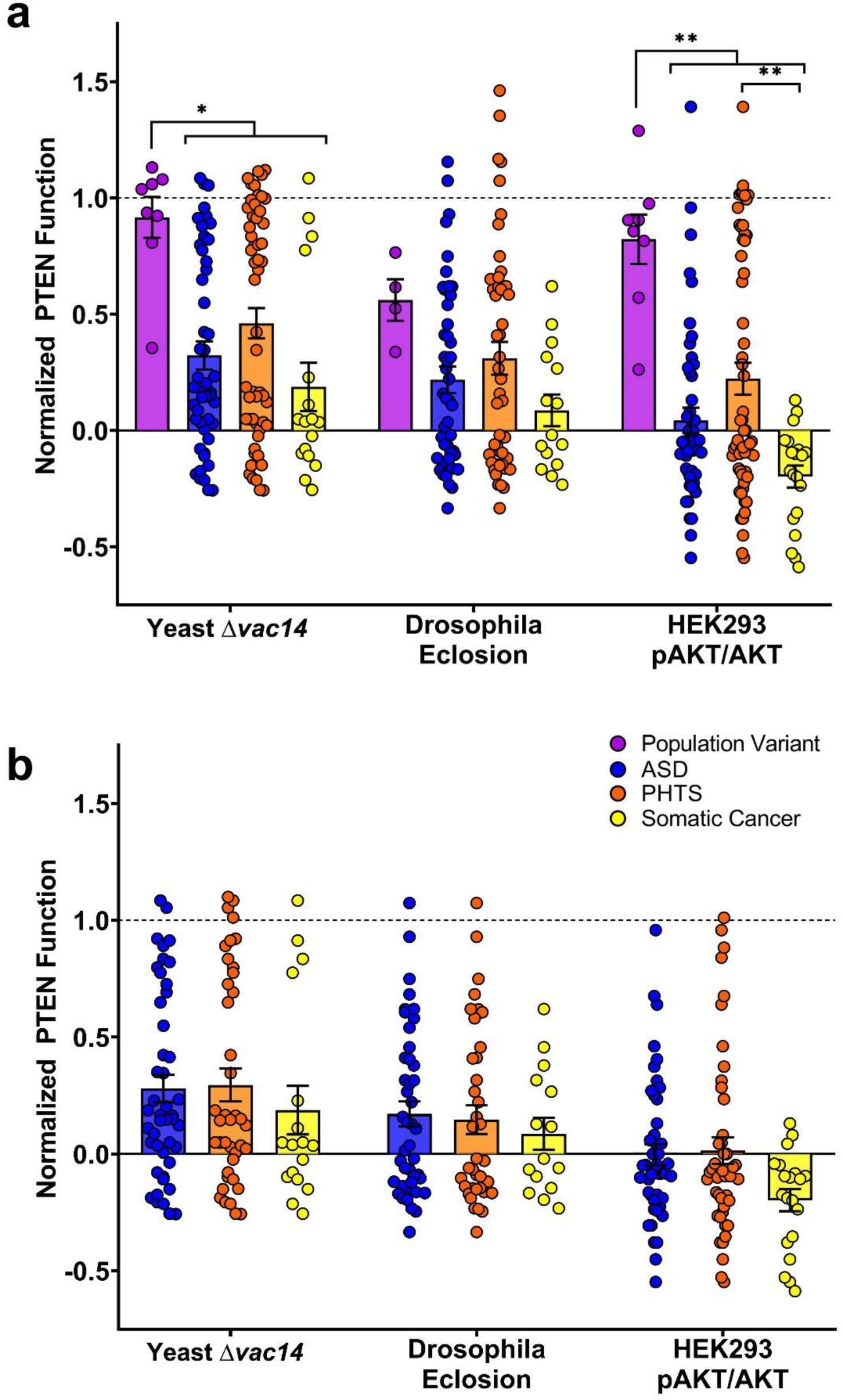
Distinct molecular mechanisms of PTEN dysfunction cannot predict ASD, PHTS or Somatic Cancer due to co-occurrence of the same variants in multiple diseases. **a** Distribution of normalized PTEN variant function by associated disease phenotypes in three high-throughput functional assays. Variants within each phenotype subgroup are not mutually exclusive as several have been found in multiple syndromes/conditions (see **Fig. 6a**). **b** Data as in (**a**), removing variants we predict to be benign (**Fig. 8b**). Mean PTEN function is plotted as bars ± SEM. * p<0.05, ** p<0.005, *** p<0.0005 by two-way ANOVA.

**Supplemental Figure 7.**
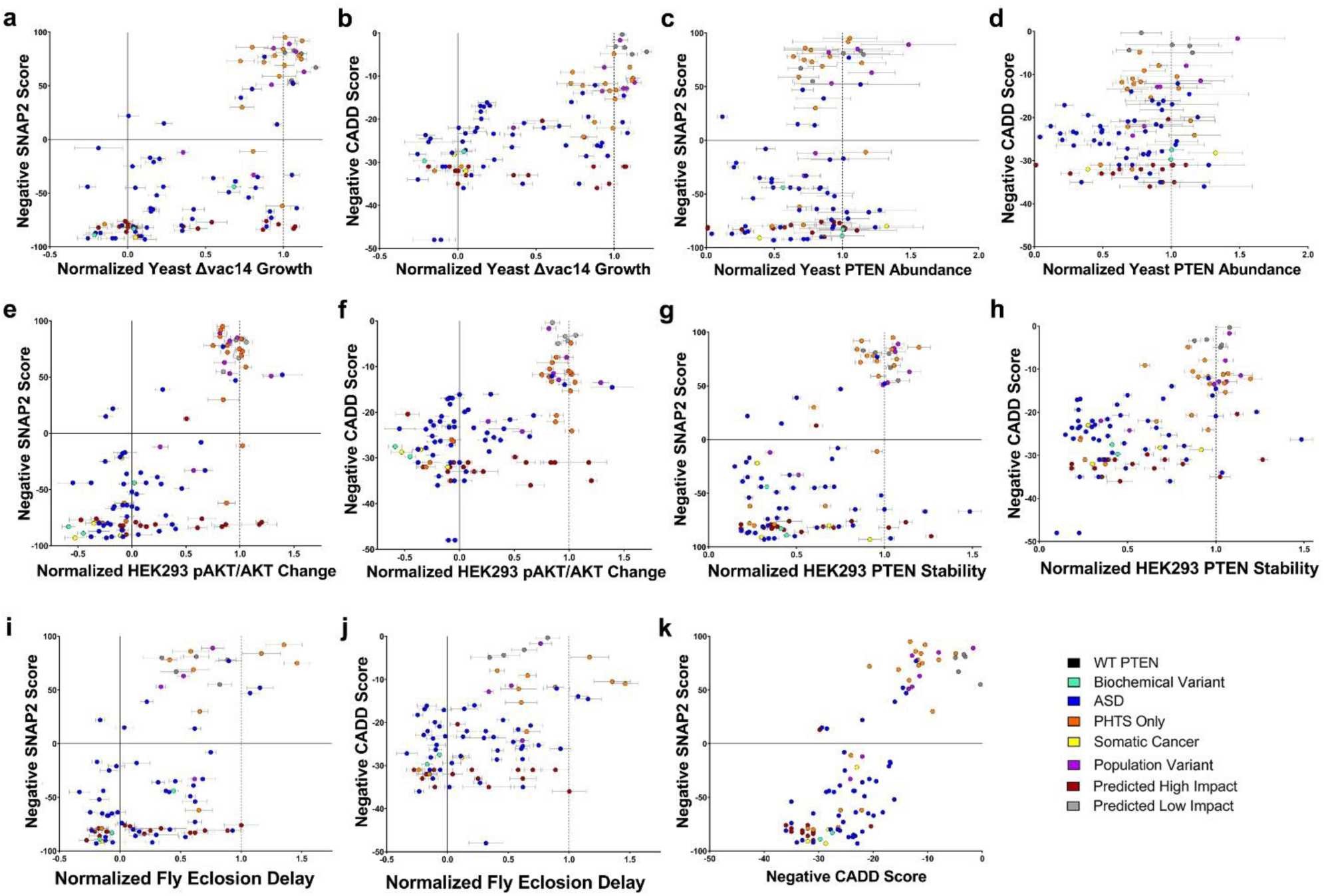
Plots of functional scores in the yeast Δ*vac14* (**a,b)** HEK293 pAKT/AKT (**e,f**), and *Drosophila* eclosion delay (**i,j**) assays compared to SNAP2 and CADD bioinformatics software predicted functional impact. Plots of PTEN variant protein abundance in yeast (**c,d**), and stability in HEK293 (**g,h**) assays compared to SNAP2 and CADD scores. **k** Plot of predicted PTEN variant functional impact by SNAP2 vs. CADD. For all plots, negative SNAP2 and CADD scores are shown so higher values predict more function, and lower values predict LoF. Pearson correlation r- and p-values for all plots presented in **Supp. Table 3**.

**SUPPLEMENTAL TABLE 1. Annotation of PTEN variants.**

**SUPPLEMENTAL TABLE 2. Mean and SEM vales for each variant testing in each assay.**

**SUPPLEMENTAL TABLE 3. Pearson coefficient r and p-values for comparisons between all assays.**

**SUPPLEMENTAL TABLE 4. *C. elegans* strains created for this study.**

## METHODS

### PTEN Variant selection and annotation

We collected PTEN missense and nonsense base substitution variants from VariCarta, SFARI Gene, ClinVar, COSMIC, and ExAC databases and the literature. We used VariCarta to harmonize and annotate the ASD-associated variants. **Biochemical Variants** were selected from the literature. **Population Variants** were selected which have relatively high allele frequency in ExAC and/or gnomAD and the absence of disease-association with CADD < 25. **Predicted High Impact** or **Predicted Low Impact** variants were selected based on the CADD phred version 1.0 31 or SNAP2 scores. SNAP2 scores were obtained by correspondence with the authors^1^. Variants were obtained in an HGVS protein mutation format and back-converted to genomic coordinates using TransVar^2^, using “Reverse Annotation: Protein” mode, GRCH37/hg19 as the reference genome and RefSeq for the annotation database. Resulting genomic coordinates were then annotated using both a local Annovar instance^3^ and the wAnnovar web server (ran on June 24th 2019) for additional annotations.

### Gene interaction screen in *Saccharomyces cerevisiae*

The synthetic dosage lethality (SDL) screen was performed using media and techniques as previously described^4,5^. The Y*7093* strain was transformed with the p*GAL1/PTEN* plasmid mated to the haploid deletion mutant array at a density of 1536 spots per plate using a Singer RoToR HDA robot (Singer Instruments, Somerset, United Kingdom). The resulting diploids were copied in triplicate onto enriched sporulation medium and incubated at 25°C for 14 days. *MATa* haploid cells were generated by germination on SC-His/Arg/Lys +Canavanine/Thiamine. Triple mutants were selected by two rounds of incubation on SC-His/Arg/Lys/Ura +Canavanine/Thiamine/G418 medium. A control set of single mutants was generated by two rounds of incubation on SC-His/Arg/Lys +Can/Thia/G418+ 5-Fluoroorotic acid. Colony size was determined under PTEN-inducing conditions with 2% galactose and 2% rafinose and ratios determined by comparing colony size on control (+Ura) versus experimental (-Ura) plates using the analysis software Balony^5^. ***Mini-Array analysis in Saccharomyces cerevisiae***: A novel sentinel deletion plate was created with the eight sentinels such that each sentinel was represented in 4×4 colonies 12 times on 1536 dense plates. This was mated with a set of master query plates which each contained seven variants and pEGH with rows of *PTEN* in an alternating pattern such that each variant or pEGH spot was paired with a PTEN spot directly beneath. In similarity with the SDL screens, the query and sentinel plates were mated then underwent sporulation under enriched sporulation conditions at 25℃ for 14 days. Haploid cells expressing the query plasmids in the deletion strains were selected for as in the traditional SDL and colony size was scored using Balony software after induction of gene expression using 2% galactose and 2% raffinose in the media.

### Developmental rate assay in transgenic *Drosophila*

Human PTEN variants were transferred into the pGW-HA.attB destination vector (PMID 23637332) by Gateway-mediated recombination. These were integrated into the *attP2* locus of the *Drosophila* genome, by phiC31-integrase (performed by Rainbow Transgenics Inc, CA) which inserts a single copy of the gene and ensures reproducible transcript expression between all variants (PMID 17360644). To measure eclosion, homozygous *da-GAL4* virgin females were crossed to heterozygous *UAS-PTEN variants/TM3, Ser, Sb,GMR-Dfd-EYFP* (TDGY) males at a ratio of 2:1. These adults are predicted to produce progeny of genotype *da-GAL4/UAS-PTEN* and *da-GAL4/TDGY* at a 50:50 Mendelian ratio. In this way, the *da-GAL4/TDGY* flies serve as an internal genetic control (that is the same in every vial examined) to which the developmental rate of *da-GAL4/UAS-PTEN* variants can be compared. For every PTEN variant tested, 3 vials of mating flies were set up and combined time to eclosion rates was measured between vials for both PTEN and TDGY progeny. For every batch of PTEN variants that were tested in parallel at the same time, we also set up a 3 vials each of *da-GAL4* crossed to WT/TDGY, attP2/TDGY and C124S/TDGY. Mating flies were maintained on standard cornmeal food at 25°C and 70% humidity at a constant 12h light/dark cycle. Prior to the assay, mating flies were housed for a 36-48h acclimatization period. For the assay, mating flies were placed on fresh yeast-augmented food for 1h to clear out older fertilized eggs, then placed in a fresh vial in which females laid freshly fertilized eggs within a 6h window. Adult flies were removed from this vial and progeny were monitored until eclosion. The appearance of the first adult fly in each vial was considered as time zero, and adult fly eclosion was assessed thereafter every 12 hours for the next 3 days, comparing the time to eclosion for EYFP-positive and EYFP-negative flies per vial.

### Neuronal morphogenesis and synaptogenesis in rat primary neuronal cultures

All experimental procedures and housing conditions were approved by the UBC Animal Care Committee and were in accordance with the Canadian Council on Animal Care (CCAC) guidelines. Dissociated hippocampi and dorsal root ganglia (DRG) were isolated from embryonic day 18.5 (E18) of Sprague-Dawley rats (Charles River, Sherbrooke, Canada) of either sex prepared as previous described^6^ and plated at a density of 130 cells/mm^2^ for hippocampus, and 50K per coverslip pre-coated with poly-D-lysine for DRG. Neurons were co-transfected with eGFP and *PTEN* variants with Lipofectamine 2000 (Invitrogen/Life Technologies, CA, USA) at 9-11 DIV for hippocampus, and 1 DIV for DRG. Hippocampal cultures were fixed at 13-14 DIV for measures of synaptogenesis and morphogenesis, and DRG were fixed at 24h after transfection for measures of axonal growth. ***Immunocytochemistry:*** Immunocytochemistry was performed as previously reported^7^. Cells were incubated with primary antibodies (1:500 mouse anti-PSD-95 (ABR, IL, USA) or 1:300 mouse anti-gephyrin (Synaptic Systems, Goettingen, Germany) overnight at 4°C in 1% goat serum/PBS, and incubated with Alexa488 or Alexa568 conjugated goat anti-mouse secondary antibodies (1:1000, Molecular Probes, ON, Canada) for 1 hour at RT. Neurons were mounted on microscope slides with Prolong Gold (Molecular Probes, Thermo Fisher Scientific, MA, USA) and imaged using an Olympus Fluoview 1000 confocal microscope. For DRG, antibodies used were Anti-Beta-Tubulin III Antibody Clone TUJ1 (STEMCELL Technologies, BC, Canada) and Texas Red dye-conjugated AffiniPure Goat Anti-Mouse IgG (Jackson ImmunoResearch Laboratories Inc., PA, USA), mounted with VECTASHIELD® Antifade Mounting Media. ***Measurement of total dendritic length:*** Dendritic morphology and soma size were imaged using a 20X/0.75 Oil Plan-Apochromat objective. ***Measurement of synaptic density and total synapses***: To quantify synaptic density, fixed neurons immunostained with antibodies against Gephyrin (inhibitory synapses) and PSD-95 (excitatory synapses). Synapse were imaged using a 60X/1.42 Oil Plan-Apochromat objective. Synaptic puncta were defined as 0.05-3 µm, and synaptic density was calculated by obtaining the total number of puncta within the dendrite divided by the total dendrite length. Puncta numbers were obtained using a custom macro that made use of the “Analyze Particles” function of ImageJ and the “Colocalization” plugin (https://imagej.nih.gov/ij/plugins/colocalization.html), and total length was measured using ImageJ^8^. ***Data analysis of total axonal length in DRG:*** Images of neurons were obtained using a Zeiss Axioplan II fluorescence microscope equipped with a Plan-NEOFluar 20x/0.50 objective lens. For each culture, 10 GFP-positive neurons were randomly sampled. To evaluate neurite length and branching, neurites (≥2µm) were traced using ImageJ 1.52a software equipped with the NeuronJ plugin.

### *C. elegans* chemotaxis assay

Worms were cultured on Nematode Growth Medium (NGM) seeded with *Escherichia coli* (OP50) as described previously^9^. N2 Bristol, and CB1375 *daf-18(e1375)* strains were obtained from the *Caenorhabditis* Genetics Center (University of Minnesota, USA). All *PTEN* variants were expressed as pan-neuronal extrachromosomal arrays injected into the *daf-18(e1375)* reduction-of-function background at 50ng/ul final DNA concentration. *daf-18(e1275)* harbors a 30–base pair insertion in the fourth exon and is predicted to insert six amino acids before introducing an early stop codon that truncates the C-terminal half of the protein while leaving the phosphatase domain intact^10^. The strains created for this work are listed in **Supp. Table 4**. ***Strain and plasmid generation:*** The *PTEN* entry clones were recombined with a pDEST-*aex-3p* destination vector for pan-neuronal expression (obtained from Dr. Hidehito Kuroyanagi^11^) to generate the *aex-3p::PTEN::unc-54 UTR* rescue construct using gateway cloning (Invitrogen), according to manufacturers instructions. Multiple transgenic strains were generated per genotype using standard microinjection and fluorescent screening for a *pmyo-2::mCherry::unc-54 3’ UTR* (pCFJ90) cotransformation marker (3 ng/ul). ***Genotype confirmation:*** *PTEN* variant sequences were confirmed by amplifying the entire CDS PCR followed by Sanger sequencing. The forward and reverse primers used to amplify the *PTEN* CDS were ATGACAGCCATCATCAAAGA and TCAGACTTTTGTAATTTGTG respectively. ***Multi-Worm Tracker NaCl chemotaxis behavioral assays:*** The chemotaxis behavioral assay was conducted as previously described^12,13^. Briefly, assays were conducted on a 6 cm assay plate (2% agar) where a salt gradient was formed overnight by inserting a 2% agar plug containing 50mM of NaCl (approximately 5 mm in diameter) 1 cm from the edge of the plate. A control 2% agar plug without NaCl was inserted 1 cm from the opposite edge of the plate. Strains were grown on NGM plates seeded with *E. coli* (OP50) for 3 or 4 days. Worms on the plates were collected and washed three times using M9 buffer before being pipetted onto an unseeded NGM plate to remove excess buffer and select animals carrying transformation markers. Adult worms were transferred and placed at the centre of the assay plates and tracked for 40 minutes on the Multi-Worm Tracker^14^. After the tracking period, the chemotaxis index was calculated as (A – B)/(A + B), where A was the number of animals that were located in a 1.5 cm-wide region on the side of the assay plate containing the 2% agar plug with 50mM NaCl and B was the number of animals that were located in a 1.5 cm-wide region on the side of the assay plate containing the 2% agar plug without NaCl. Animals not located in either region (ie. the middle section of the assay plate) were not counted towards the chemotaxis index. One hundred to two hundred animals were used per plate, and two or three plate replicates were used for each line in each experiment. Multi-Worm Tracker software (version 1.2.0.2) was used for image acquisition^14^. Behavioral quantification with Choreography software (version 1.3.0_r103552) used “--shadowless”, “--minimum-move-body 2”, and “-- minimum-time 20” filters to restrict the analysis to animals that moved at least 2 body lengths and were tracked for at least 20 s. Custom R scripts organized and summarized Choreography output files. Each experiment was independently replicated at least twice. No blinding was necessary because the Multi-Worm Tracker scores behavior objectively.

### Variant protein abundance measures in *Saccharomyces cerevisiae*

Yeast were grown overnight under inducing conditions. A_600_ was measured for all samples then each were diluted to an OD of 0.1 and allowed to grow approximately 6-8 hours or until they reached between 0.8-1.0OD. An equivalent of 1.0 OD was collected for each sample. Samples were pelleted and supernatant discarded before being frozen at −20C. After freezing, glass beads and sample buffer containing sodium dodecyl sulfate (SDS) and 2.0% β-mercaptoethanol were added to the pellet then bead bashed for 2 minutes. Samples were then boiled at 90C for 5 minutes and centrifuged before being loaded into a 4-12% Bis-Tris protein gel (Thermo Fisher Scientific, cat #NP0322) and run with MOPS running buffer (Thermo Fisher Scientific, cat #NP0001). Gels were transferred in buffer containing 20% methanol and 0.03% SDS for 2 hours at 4C at 28V onto 0.2μM nitrocellulose membranes (Bio-Rad, cat #1620097). Membranes were blocked in tris-buffered saline and Tween 20 (TBST) (Acros Organics, cat #AC233360010) containing 5% milk (Bio Basic, cat #NB0669). Membranes were then stained with PTEN antibody (1:1000) (R&D Systems, cat# MAB847,) followed by goat-anti-mouse HRP (Thermo Fisher Scientific, cat#62-6520) and imaged with ChemiDoc MP (Bio-Rad). Membranes were then stripped with a pH 2.2 stripping buffer containing 1.5% glycine, 0.1%SDS and 1% Tween 20 and re-stained for the loading using the anti-glucose-6-phosphate dehydrogenase antibody (Millipore, cat #A9521). Blots were again imaged on ChemiDoc MP (Bio-Rad). Band density was determined using ImageJ. All PTEN bands were normalized first to their loading control then the wildtype PTEN done in each experiment. All variants were tested in triplicate.

### Immortalized Cell Culture, Protein stability and pAKT/AKT assays

HEK293 cells purchased from the American Type Culture Collection (CRL-1573) and were routinely passaged in Dulbecco’s Modified Eagle’s Medium (DMEM) (MilliporeSigma D6046) supplemented with 10% FBS and 100U/mL Penicillin-Streptomycin. For all experiments herein, HEK293 cells were used for a maximum of 15 passages. Cells were seeded at 1×10^5^ per well in 24 well dishes 16-20hrs before transfected with 500ng of expression plasmid using X-tremeGENE 9 at a ratio of 2uL to 1ug DNA. 24h after transfection cells were washed with DMEM and starved for 19h in serum-free media before being stimulated for 10 minutes with complete medium plus 10nM Insulin (12585014, Gibco), then washed once in 1xPBS before treated with Trypsin-EDTA (25200072, Gibco) for 5 minutes to create a single-cell suspension and then fixed for 10 minutes in 3.2% PFA. Cells were permeabilized with Flow Cytometry Permeabilization/Wash Buffer I (FC005, R&D Systems) and stained with 1:100 of Rabbit anti-pAKT (S473, #9271, Cell Signaling) and 1:100 of Mouse anti-pan-AKT (#2920, Cell Signaling) for one hour on ice. Cells were washed and then stained with 1:100 of Goat anti-Rabbit IgG-Alexa Fluor 647 (A21244, Thermo Fisher) and 1:100 of Goat anti-Mouse IgG-Alexa Fluor 405 (A31553, Thermo Fisher) for one hour on ice. Cells were washed and re-suspended in Flow Cytometry Staining Buffer (FC001, R&D Systems) before loading into an Attune Nxt Flow Cytometer (Invitrogen)***. Flow Cytometry Analysis:*** Data was recorded using VL-1 (Alexa Fluor 405), BL-1 (sfGFP), YL-1 (mtagRFP-T) and RL-1 (Alexa Fluor 647) channels, which were single-stain compensated. Using FlowJo, Cells were selected using FSC-H/SSC-H and single cells were selected using SSC-H/SSC-A. For pAKT/AKT measures, cells with sfGFP fluorescence above untransfected cells to 100-fold above untransfected (300 a.u. to 20’000 a.u.) were selected. For stability measures, cells with higher mtagRFP-T fluorescence than untransfected and sfGFP fluorescence <20’000 a.u. were selected. The median of (647 - Background) / (405 -Background) as the pAKT/AKT level was calculated both for the positive and negative (RFP- & sfGFP-, untransfected) population in each sample and the difference was measured as the effect measure, normalizing each sample by its in-well untransfected control. Samples were then normalized to WT=1 for each day of experiment. For stability measures, the median ratio of (sfGFP - Background)/(mtagRFP-T – Background) was calculated as the relative stability of PTEN compared to its 1:1 transfection control. Samples were then normalized to WT=1 for each day of experiment.

### Data modeling and variant effects analysis

The quantitative phenotypes for the yeast, fly, worm and rat assays were analyzed using a hierarchical (mixed effect) model approach to account for experimental batch effects and variability between replicates. For each assay, we treat the genotype as a fixed effect, which can be either the positive control, the negative control, or a variant, and treat blocking factors such as experimental batch or day as random effects (stratified by variant and accompanying controls). All R scripts are provided at https://github.com/PavlidisLab/Post-PTEN. For each assay, we fit models to the data for all tested variants and accompanying controls jointly. Given a genotype *i* and a sample *j*, we fit:

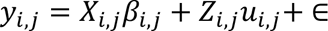

Where y is the observed quantitative phenotype (length *N* where *N* is the number of data points for the assay in total), *X* and *Z* are the model matrices for fixed and random effects respectively, is a vector of parameters for the fixed effects (the genotypes of each variant), u is vector of parameters for random effects, and is residual error. Models were fit using the lmer function of the lmerTest package^15^ in the R programming environment <u>(R Core Team 2019)</u>. To assess the significance of effect between the positive (wild type PTEN) and negative (no PTEN, empty vector or GFP background) controls, we extracted the model fit p-values of each genotype under a comparison where each control is used as the contrast group. Both comparison yields a vector of p-values for all other genotypes except the contrast variant. For visualization purposes, we plot data adjusted for the estimated random effects. Specifically, we compute adjusted values through a multiplication between the fixed-effects model matrix X with the β fixed-effect parameter estimate and a sum of the model residuals vector. In other words:

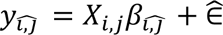

We then take the y values for each genotype and rescale the data such that the mean positive control equals 1.0, and the mean negative control equals 0.0. Finally, we rescale the standard errors for each genotype from the adjusted data proportionally to the ratio of the adjusted data and the normalized data distributions, thus keeping the values consistent with the 0.0-1.0 re-scaling. Graphical Analysis. All results are presented as means ± SEM. For HEK293 experiments, data were analysed with Pearson correlations tests and Student’s t-tests with WT unless otherwise indicated. All statistical analysis was done using the statistical programming software Graphpad p ≤ 0.05 was considered as significant.

