## Supplementary material for "Multi-model functionalization of disease-associated *PTEN* missense mutations identifies multiple molecular mechanisms underlying protein dysfunction": C elegans strains used

**SUPPLEMENTARY TABLE 4. *C. elegans* strains created for this study.**

VG667-668, 672-673 *yvEx667-668, 672-673[paex-3::PTEN::unc-54; pmyo-2::mCherry::unc-54 UTR]*

VG674, 810-816 *daf-18(e1375);* *yvEx674, 810-816[paex-3::PTEN::unc-54; pmyo-2::mCherry::unc-54 UTR]*

VG723-724 *daf-18(e1375); yvEx723-724[paex-3::PTEN*-C124S*::unc-54; pmyo-2::mCherry::unc-54 UTR]*

VG718-719 *daf-18(e1375); yvEx718-719[paex-3::PTEN*-G132D*::unc-54; pmyo-2::mCherry::unc-54 UTR]*

VG716-717 *daf-18(e1375); yvEx716-717[paex-3::PTEN*-Y176C*::unc-54; pmyo-2::mCherry::unc-54 UTR]*

VG720-721,722-723 *daf-18(e1375); yvEx720-721[paex-3::PTEN*-H123Q*::unc-54; pmyo-2::mCherry::unc-54 UTR]*

VG718-719 *daf-18(e1375); yvEx718-719[paex-3::PTEN*-G132D*::unc-54; pmyo-2::mCherry::unc-54 UTR]*

VG725-726 *daf-18(e1375); yvEx725-726[paex-3::PTEN-I101T::unc-54 UTR; pmyo-2::mCherry::unc-54 UTR]*

VG727, 762-764 *daf-18(e1375); yvEx727, 762-764[paex-3::PTEN-D326N::unc-54 UTR; pmyo-2::mCherry::unc-54 UTR]*

VG729-731 *daf-18(e1375); yvEx729-731[paex-3::PTEN-D92N::unc-54 UTR; pmyo-2::mCherry::unc-54 UTR]*

VG732-733, VG752-753 *daf-18(e1375); yvEx732-733, 752-753[paex-3::PTEN-R130L::unc-54 UTR; pmyo-2::mCherry::unc-54 UTR]*

VG754, 756 *daf-18(e1375); yvEx754-756[paex-3::PTEN-P38H::unc-54 UTR; pmyo-2::mCherry::unc-54 UTR]*

VG758-761 *daf-18(e1375); yvEx758-761[paex-3::PTEN-T167N::unc-54 UTR; pmyo-2::mCherry::unc-54 UTR]*

VG765-768 *daf-18(e1375); yvEx765-768[paex-3::PTEN-T131I::unc-54 UTR; pmyo-2::mCherry::unc-54 UTR]*

VG769-772 *daf-18(e1375); yvEx69-772[paex-3::PTEN-H93R::unc-54 UTR; pmyo-2::mCherry::unc-54 UTR]*

VG773-776 *daf-18(e1375); yvEx773-776[paex-3::PTEN-D268E::unc-54 UTR; pmyo-2::mCherry::unc-54 UTR]*

VG777-779 *daf-18(e1375); yvEx777-779[paex-3::PTEN-G44D::unc-54 UTR; pmyo-2::mCherry::unc-54 UTR]*

VG780-785 *daf-18(e1375); yvEx780-785[paex-3::PTEN-Q171E::unc-54 UTR; pmyo-2::mCherry::unc-54 UTR]*

VG792-795 *daf-18(e1375); yvEx792-795[paex-3::PTEN-K6E::unc-54 UTR; pmyo-2::mCherry::unc-54 UTR]*

VG796-798 *daf-18(e1375); yvEx796-798[paex-3::PTEN-A79T::unc-54 UTR; pmyo-2::mCherry::unc-54 UTR]*

VG805-806 *daf-18(e1375); yvEx805-806[paex-3::PTEN-K6I::unc-54 UTR; pmyo-2::mCherry::unc-54 UTR]*

VG807-809 *daf-18(e1375); yvEx807-809[paex-3::PTEN-P354Q::unc-54 UTR; pmyo-2::mCherry::unc-54 UTR]*
